# Combined Menin and XPO1 inhibition drive synergistic antileukemic activity in *KMT2A*r and *NPM1*-m AML

**DOI:** 10.64898/2026.03.10.710924

**Authors:** Md. Hafiz Uddin, Sandhya Dhiman, Yufen Han, Amro Aboukameel, Vikram Dhillon, Jeff Aguillar, Steven Buck, Abhinav Deol, Julie L Boerner, Lisa Polin, Linda Kessler, Francis Burrows, Jay Yang, Asfar S. Azmi, Jaroslaw P. Maciejewski, Jevon Cutler, Yang Du, Suresh Kumar Balasubramanian

## Abstract

Menin scaffolds the oncogenic histone-lysine-N-methyltransferase (KMT2A)-fusion protein (FP) complex in *KMT2A-*r and wild-type *KMT2A* complex in *NPM1*-m acute myeloid leukemia (AML). Menin inhibitors (MIs) are effective in *KMT2A*-r AML and *NPM1*-m AML. However, not all patients respond to MIs as monotherapy. In this preclinical study, we demonstrate that the MI ziftomenib, in combination with the XPO1 inhibitor selinexor, synergistically inhibited the growth of multiple *KMT2A-*r and *NPM1*-m AML cell lines (CI<1). The combination suppressed colony formation in primary CD34+ *KMT2A-*r progenitor cells without affecting normal stem cells. Robust apoptosis and decreased G2/M populations were also evident. The combination downregulated HOXA9 and MEIS1 while upregulating monocytic differentiation marker CD11b in both the AML molecular signatures. RNA sequencing and proteomic analysis in *KMT2A-*r revealed suppression of multiple bona fide menin-KMT2A target genes. Our mechanistic studies also identified a novel role of XPO1 in stabilizing menin’s binding to chromatin and its interactions with KMT2A and KMT2A/MLLT3. XPO1 inhibitor-mediated disruption of these interactions, particularly in combination with ziftomenib, synergistically impairs oncogenic transcriptional programs. *In vivo*, combination therapy improved survival in both MV4;11 and OCI-AML3 cell line and primary patient-derived *KMT2A*-*r* and *NPM1*-m AML xenograft models in NSG mice, effective even at reduced drug doses. These preclinical findings demonstrate that simultaneous inhibition of the menin-KMT2A interaction and XPO1 can be a more effective translational strategy for treating *KMT2A-*r and *NPM1*-m AML than MI monotherapy to deepen responses and delay/prevent relapses.

## Introduction

The *MEN1* gene, located on chromosome 11q13, encodes menin, a scaffold protein essential for coordinating protein-protein interactions that organize critical cellular processes, including gene transcription, DNA repair, cell cycle control, and apoptosis^1, 2^. Menin binds to the menin-binding domain (MBD) of histone-lysine-N-methyltransferase/Mixed lineage leukemia1 (KMT2A/MLL1), anchoring it to chromatin and facilitating the transcription of key leukemogenic targets in *KMT2A*-rearranged (r) acute myeloid leukemia (AML)^3, 4^. In *KMT2A*-r AML, chromosomal translocations involving *KMT2A* generate over 80 fusion proteins (FP) that retain the interaction with menin and aberrantly activate downstream genes, notably *HOXA9* and its cofactor *MEIS1*, to drive malignant transformation^5^. Intriguingly, this same *HOX*/*MEIS1*-driven leukemogenic transcriptional program is also activated in *NPM1*-mutant (*NPM1*-m) AML, accounting for over 30% of newly diagnosed adult AML patients despite lacking *KMT2A*-r^6, 7^. In *NPM1*-m AML, NPM1c aberrantly associates with specific regions in the chromatin and cooperates with the WT KMT2A-menin complex to sustain leukemogenic gene expression, particularly via RNA polymerase II engagement and *HOX*/*MEIS1* gene activation, identifying menin-KMT2A complex as a critical target for the treatment of this AML subtype.

Menin inhibitors (MIs), developed to disrupt the oncogenic menin-KMT2A FP association, were initially intended to target *KMT2A*-r leukemias^8^; their efficacy was later extended to *NPM1*-m AML due to shared transcriptional dependencies^9^. By evicting menin from chromatin, these inhibitors downregulate leukemogenic transcription and restore myeloid differentiation^10^. Their introduction has redefined the therapeutic space for AML by offering a potent, primarily non-cytotoxic treatment tool for genetically defined subgroups. Several MIs continue to advance through clinical development (NCT04067336, NCT05735184, NCT06001788, NCT05886049, NCT05761171, NCT04811560). Revumenib has recently received FDA approval for relapsed or refractory (r/r) *KMT2A*-r and *NPM1*-m AML, and ziftomenib is now approved for r/r *NPM1*-m AML based on the encouraging safety and efficacy signals in early-phase trials across *KMT2A*-r and *NPM1*-m AMLs^11–13^. As with many monotherapeutic approaches in AML, a number of patients may not respond primarily, and responses among initial responders could be short lived. While mechanisms of acquired resistance have been described in revumenib-treated patients and supported by *in vitro* models, resistance pathways to other MI remain incompletely defined and warrant further investigation^14, 15^.

Rational combination therapies enhance efficacy while allowing dose reduction and mitigating resistance, are often considered superior to monotherapy in many hematological malignancies. To this end, we examined targeted inhibition of XPO1 in combination with MI. Nuclear export is predominantly performed by exportin 1 (XPO1, also known as CRM1), a karyopherin that exports numerous leucine-rich nuclear export signal (NES)-bearing cargo proteins from the nucleus to the cytoplasm. In many cancers, including AML, XPO1 is dysregulated or overexpressed, leading to cytoplasmic mislocalization and functional inactivation of vital tumor suppressors including p53, p21, p27 and Rb. Pharmacologic inhibition of XPO1 retains the key proteins and restores its tumor suppressor function in the nucleus^16^. Nuclear export inhibition has shown robust anti-leukemia activity in numerous preclinical and clinical studies with distinct molecular signatures^17–21^. In addition, increasing evidence has suggested that XPO1 may directly regulate transcription. XPO1 has been found to interact with transcriptional regulators including CALM/AF9, NUP98/HOXA9, SETBP1, SET/NUP214, and NPM1c and help recruiting them to target chromatin^22–27^. MIs and XPO1 inhibitors possess both independent and cooperative mechanisms of antitumor activity in *KMT2A*-r and *NPM1*-m AML. For instance, MIs can activate a tumor-suppressive program in *KMT2A*-r leukemia independently of *HOX*/*MEIS1* downregulation, via a non-canonical transcriptional program through histone demethylase UTX, to activate differentiation and cell cycle checkpoint genes^28^. XPO1 inhibition also ameliorates *NPM1*-m AML by restoring the interaction of NPM1c/PU.1 complex in the nucleus with CEBPA and RUNX1, thereby reactivating a number of granulomonocytic differentiation genes. Where the two pathways converge, XPO1 inhibitors also impair the ability of NPM1c to sustain *HOX*-driven leukemic transcription^27^ by inhibiting its recruitment to the *HOX* cluster region via chromatin-based XPO1^26^.

Hence, in the present study, we hypothesized that menin and XPO1 inhibition would synergistically suppress *KMT2A*-r and *NPM1*-m AML cell proliferation and survival. We used the clinical-stage MI ziftomenib and XPO1 inhibitor selinexor to target menin-KMT2A protein-protein interactions and XPO1 simultaneously. We also explored the role of XPO1/menin interaction in *KMT2A*-r AML.

## Materials and methods

### Cell lines, reagents, and antibodies

MV4;11 and MOLM13 cell lines were purchased from the American Type Culture Collection (ATCC, Manassas, VA, USA). OCI-AML3 and SEM cell lines were purchased from Deutsche Sammlung von Mikroorganismen und Zellkulturen (DSMZ)-German Collection of Microorganisms and Cell Cultures GmbH. The IMS-M2 cell line was kindly provided by Dr. Yogen Saunthararajah (Cleveland Clinic, USA). MOLM13 and IMS-M2 cells were maintained in RPMI-1640 medium, OCI-AML3 were cultured in MEM-α whereas MV4;11 and SEM cells were maintained in Iscove’s Modified Dulbecco’s Medium (IMDM). The RPMI-1640 medium was supplemented with 15% fetal bovine serum (FBS) (Sigma-Aldrich; catalog no. F0926) and IMDM with 10% FBS. All media were supplemented with 100 U/mL penicillin and 100 μg/mL streptomycin (GE Healthcare; catalog no. SV30010). The cell lines were maintained in a humidified incubator with 5% CO_2_ atmosphere at 37°C. The cell lines were authenticated in a core facility of the Applied Genomics Technology Center at Wayne State University [MV4;11 and OCI AML3: 04/04/2025, IMS-M2 09/02/2024, MOLM13: 09/02/2024]. The method used for testing was a short tandem repeat (STR) profiling using the PowerPlex® 16 System from Promega (Madison, WI, USA). Ziftomenib (supplied by Kura Oncology, San Diego, CA) and selinexor (supplied by Karyopharm Therapeutics, Newton, MA) were dissolved in dimethyl sulfoxide (DMSO). MI-3454 was purchased from MedChemExpress (catalog no. HY-136360). Primary antibodies used for western blot analysis were menin polyclonal antibody (Invitrogen catalog no. PA5-83140); anti-HOXA9 rabbit polyclonal antibody (Biorad; catalog no. VPA00627); Antibodies used that were purchased from Cell Signaling Technology were HOXA10 (CST, catalog no. 58891), MEIS1 rabbit antibody (catalog no. 12744S), Cyclin D1 (E3P5S) rabbit monoclonal antibody (catalog no. 55506S), CD11b/ITGAM (D6X1N) rabbit monoclonal antibody (catalog no.49420), and GAPDH (14C10) rabbit monoclonal antibody (catalog no. 2118S). Anti-β-ACTIN (Sigma; catalog no. A2228). The secondary antibodies used were IRDye 680RD Goat anti-rabbit IgG secondary antibody (LICOR; catalog no. 926-68071) and IRDye 800CW Goat anti-mouse IgG secondary antibody (LICOR; catalog no. 926-32210).

### Cell viability analysis

A total of 5000 MV4;11 cells were plated in each well of a 96-well plate. After plating, the cells were treated with either DMSO or ziftomenib (5.6 nM to 180 nM) and/or selinexor (9.3 nM to 300 nM). MOLM13 cells were seeded and treated with either DMSO or ziftomenib (36.2 nM to 1160 nM) and/or selinexor (7.5 nM to 240 nM). Similarly, OCI-AML3 were seeded and treated with either DMSO or ziftomenib (2.8 nM to 2000 nM) and/or selinexor (4.8 nM to 5000 nM). IMS-M2 were also seeded and treated with either DMSO or ziftomenib (39 nM to 5000 nM) and/or selinexor (4 nM to 5000 nM). The cells were treated with a combination of both inhibitors in a fixed ratio. The cells were incubated at 37°C and 5% CO_2_ in a humidified incubator (PHCbi model no. MCO-170AICUVHL-PA) for 72h following treatment. Cell viability was analyzed using Cell Titer-Glo (Promega; Catalogue no. G924B) following the manufacturer’s instructions. The cell viability was computed using a Synergy H1 microreader plate (Biotek). The dose-response of the drug was analyzed using GraphPad Prism (v 8.1). IC_50_ values were calculated for drugs using GraphPad Prism Software (GraphPad Software, San Diego, CA, USA). The combination index (CI) was calculated, and isobolograms were generated using Calcusyn 2.1 software (Biosoft, Cambridge, UK).

### Apoptosis assay

The apoptotic effect of inhibitors was determined using Annexin V-FITC apoptosis detection kit (Beckman Coulter Life Sciences catalog no. IM3546) according to the manufacturer’s protocol. MV4;11 cells treated with ziftomenib (5.62 nM to 180 nM), selinexor (9.37 nM to 300 nM) and their combination for 72 hours and OCI-AML3 treated with ziftomenib (2 µM), selinexor (400 nM) and their combination for 72 hours were stained with annexin V-FITC and propidium iodide (PI). The stained cells were analyzed using a Becton Dickinson flow cytometer at the Karmanos Cancer Institute Flow Cytometry Core within 1h of staining.

### Cell Cycle Analysis

The flow cytometry (FACS) was used to analyze the arrest of the specific cell cycle phase of AML cells. MV4;11 cells (2×10^6^ cells) were seeded into six well plates with 2 mL media and incubated with ziftomenib (25 nM) and selinexor (40 nM) for 48 hours. Similarly, OCI-AML3 cells (3×10^6^ cells) were also seeded with 2mL media into six well plates and incubated with ziftomenib (200 nM) and selinexor (80 nM) for 48 hours. Cells were harvested by centrifugation and washed with ice-cold PBS. Cells were fixed in 100% ice-cold ethanol and stored at 4 °C. At the time of analysis, cells were washed with ice-cold PBS and stained with freshly made 0.1 mg/mL propidium iodide (PI), 0.1% Triton x-100, and 0.2 mg/mL ribonuclease A in PBS for 30 min at room temperature in the dark. The cell cycle was then analyzed by Becton Dickinson flow cytometer at the Karmanos Cancer Institute Flow Cytometry Core.

### Colony forming unit assay

Bone marrow aspirate blood was collected from a 65-year-old male patient with *KMT2A*-r AML (MLL-3) who presented with *denovo* disease and no history of prior chemotherapy/radiation with a white cell count at diagnosis of 210,000. The patient later succumbed to primary refractory disease to multiple chemotherapies used to treat *KMT2A*-r leukemia. The CD34+ cells (HSPCs) at diagnosis were isolated by using positive selection of anti-CD34 immunomagnetic beads following the manufacturer’s protocols (EasySep™ Human CD34 Positive Selection Kit II, Stemcell Technologies; Catalog no. 17856). These CD34+ HSPC were used either freshly or after cryopreservation for further analysis. Primary CD34+ *KMT2A*-r AML cells (KCI-MLL3) were treated in T25 flasks (Ultracruz, catalog no. SC-200262) with DMSO, MI-3454 (36nm), selinexor (55nm), and the combination and viable cells were counted under a microscope (Zeiss Primovert microscope, Carl Zeiss Microscopy, Oberkochen, Germany) with trypan blue (Life Technologies, catalog no.193162) staining at the end of 12 hours. After washing, the 1000 12-hour treated and viable CD34+ primary *KMT2A-*r (MLL-3) cells were mixed with 1 ml each of MethoCult media (Stem Cell Technologies, catalog no.04100) and plated in 35mm Petri dishes (Santa Cruz Biotechnologies, catalog no. sc-351864) and kept in a 5% CO_2_ in a humidified incubator. Also additionally, 1000 primary untreated CD34+ *KMT2A*-r (MLL-3) were mixed and treated with DMSO, MI-3454 (5nm and 10nm), selinexor (9nm and 18nm), and their combinations in 1 ml each of MethoCult Express (STEMCELL), on 35 mm Petri dishes and cultured for 14 days to form colonies. Colonies were counted under the inverted phase contrast microscope using STEMgrid™-6 (Stem Cell Technologies, catalog no, 27000) at the end of 14 days in culture.

### Toxicity assay

Primary normal human CD34+ cells (Stemcell Technologies, Catalog no. 200-0001) were treated in T25 flasks with regular media (RPMI1640, Life Technologies, Catalog no.11875093) with DMSO, 36 nm of ziftomenib, 55 nm of selinexor and the combination in a 5% CO_2_ humidified incubator. The viability was assessed at 72 hours using trypan blue (Life Technologies, catalog no.193162) staining under an inverted phase contrast microscope.

### RNA extraction and RT-qPCR analysis

Total RNA was extracted from MV4;11 and OCI-AML3 cells using Qiazol lysis reagent (Qiagen; catalog no.1038703) following the manufacturer’s protocol. The concentration and purity of extracted RNA were analyzed using a spectrophotometer (Nanodrop 2000; Thermo Scientific).

The complementary DNA was synthesized from isolated RNA by reverse transcription using a cDNA synthesis kit (Applied Biosystem, Waltham, MA, USA) following the manufacturer’s protocol. cDNA was amplified using SYBR green supermix (Applied biosystem, Waltham, MA, USA) by Real-Time PCR (Applied Biosciences Quant Studio 3 real-time PCR system). The sequences of primers used for RT-qPCR (human and mouse) in this study are listed in Supplementary Table S1 and S2. The SYBR Green master mix was activated by heating at 95°C for 10 min. A total of 40 thermal cycles were used, consisting of 15 s at 95°C and 1 min at 60°C steps followed by melting curve analysis. The glyceraldehyde 3-phosphate dehydrogenase (GAPDH), and 18s rRNA were used as reference genes in this experiment. The relative gene expression was analyzed following the ΔΔCt method (Livak and Schmittegen, 2001).

### Nuclear and cytoplasmic protein extraction

Nuclear and cytoplasmic proteins were fractionated from OCI-AML3 cells using NE-PER nuclear and cytoplasmic extraction reagents following the manufacturer’s instructions. Extracted lysates were stored at −80 °C for further use (Thermo Fisher Scientific, Waltham, MA; catalog no. 78833).

### Western blot analysis

A total of 1×10^6^ MV4;11 and OCI-AML3 cells were seeded in each T25 flasks. Different doses of ziftomenib, selinexor and a combination of both drugs were added and incubated up to 72 hours. Following the incubation, total protein was extracted from the treated cells using RIPA buffer following the manufacturer’s protocol (Chem Cruz, Sc-24948A). The protein concentration was determined using Pierce™ BCA Protein Assay Kits (Thermo Fisher Scientific, catalog no. 23225). The sample was prepared with the Laemmli SDS sample buffer (Thermo Fisher Scientific, J61337AD) and heated at 95° C for 10 min on a heat block (Benchmark Scientific, USA). The protein samples were subjected to SDS-PAGE using precast gel (Bio-Rad, Hercules, CA, USA) and then transferred onto nitrocellulose (Amersham, Germany; catalog no. 10600006) or polyvinylidene fluoride (PVDF) membranes (Sigma, IPVH00010) by electrophoresis. The membranes were blocked with 5% skim milk or bovine serum albumin (BSA) (Sigma-Aldrich: catalog no. A7906) in 1X PBS (Biorad, catalog no. 1610780) with 0.05% tween (PBST) at room temperature for 1 hour and then incubated overnight with primary antibodies (1:1000) at 4°C. Membranes were washed three times with PBST following the incubation and then probed with secondary antibodies (1:10000) for one hour at room temperature. After the incubation, membranes were washed with PBST three times, and protein was visualized using Odyssey DLx Imaging System by LI-COR (Lincoln, NE, USA).

### Immunoprecipitation

HEK293T cells were transfected with the *pMSCVpuro-KMT2A/MLLT3* using LipoD293 (SignaGen Laboratories) and nuclear extracts were isolated 48 hours after transfection and 6 hours after treatment with KPT-185, selinexor, ziftomenib, or DMSO using the Nuclear Complex Co-IP kit (Active Motif). Nuclear lysates were incubated with 2 µg of anti-menin (Fortis Life Sciences, Cat. #A300-115A), anti-KMT2A-N (Fortis Life Sciences, Cat. #A300-086A), or control IgG (Cell Signaling Technology, Cat. #2729) for overnight at 4°C. Subsequently, Dynabeads protein G (Invitrogen, Carlsbad, CA) were added to the nuclear lysate/antibody mixture and incubated for an additional 2 hours at 4°C. Immunoprecipitated proteins were eluted and analyzed by Western blotting analysis. For protein detection following antibodies were used: anti-KMT2A-N (Fortis Life Sciences, Cat. #A300-086A), anti-MEN1 (Fortis Life Sciences, Cat. #A300-115A), anti-XPO1 (Cell Signaling, Cat. #46249, Abcam), anti-β-Actin (Millipore, Cat. #MAB1501R) and anti-Rabbit TrueBlot® secondary antibody (Rockland, Cat. #18-8816-31). Protein bands were visualized by incubation with SuperSignal West Femto chemiluminescent substrate (Pierce). Images were captured using ChemiDoc MP imaging system (Bio-Rad).

### Retroviral constructs and generation of retroviruses

pMSCVpuro-KMT2A/MLLT3 was generated by retrieving KMT2A/MLLT3 cDNA from pMIG-MLL-AF9 with *EcoR*I digest followed by ligation with the pMSCVpuro vector. Menin cDNA was amplified from mouse myeloid progenitor cells by RT-PCR, confirmed by sequencing, and cloned into the *BamH*I and *Not*I site of pMYs-IRES-GFP vector for the generation of *pMYs-Men1-IRES-GFP*. The primers are: Men1 S, 5’-CGC GGG ATC CGC CGC CAT GGG GCT GAA G-3’; Men1 AS, 5’-CGC GGC GGC CGC TCA GAG TTC AGA GGC CCT TG-3’. NES1 and NES2 mutations were introduced into menin cDNA using QuickChange II site-directed mutagenesis kit (Agilent Technologies, Santa Clara, CA) and the mutant cDNAs were also cloned into pMYs-IRES-GFP vector. A 3xFLAG tag was added to C-terminus of the wild-type and mutant menin proteins by PCR for assessing their cellular localization. High titer retroviruses were produced by transient transfection of Plat-E cells using Fugene-6 (Roche, Indianapolis, IN). Viral titers were determined by serial dilution and infection of NIH-3T3 cells. Retroviral transduction was carried out as previously described^24^.

For the generation of lentivirus, pLKO.5 lentiviral constructs were purchased from Sigma Aldrich (NC-sh, SHC202; Men1-sh, TRCN0000310893). The generation of infectious lentivirus and lentiviral transduction were performed as previously described^24^.

### Generation of KMT2A/MLLT3-immortalized myeloid progenitor cells

Freshly purified Lin-Sca-1+c-kit+ (LSK) cells from female C57BL/6 mice were initially cultured in DMEM media with 15% heat inactivated fetal bovine serum, and mouse cytokines (100 ng/ml SCF, 6 ng/mL IL-3, and 10 ng/mL IL-6) for 24 hours to stimulate cell proliferation, and subsequently infected twice with pMSCVpuro-KMT2A/MLLT3 retrovirus on retronectin-coated plates (Takara Bio) for 48 hours. Transduced cells were passaged in IMDM supplemented with heat inactivated 20% horse serum, 1X penicillin/streptomycin, and mouse cytokines (50 ng/ml SCF and 10 ng/mL IL-3) for 2 weeks to establish immortalized cells.

### ChIP (Chromatin Immunoprecipitation)

Chromatin immunoprecipitation (ChIP) and sequential re-ChIP were performed using the ChIP-IT Express Enzymatic Kit (Active Motif, Cat. No. 53035) and the Re-ChIP-IT Kit (Active Motif, Cat. No. 53016), following the manufacturer’s protocols with minor modifications. Briefly, cells were crosslinked with 1% formaldehyde to preserve protein-DNA interactions and quenched with glycine. Nuclei were isolated, and chromatin was enzymatically sheared to generate DNA fragments of ∼200-1500 bp. For immunoprecipitation (ChIP), chromatin was incubated with menin-specific antibody (Fortis Life Sciences, Cat. #A300-105A; Active Motif, Cat. #61005), XPO1-specific antibody (Novus Biological, Cat. #NB100-79802), or control IgG and captured using Protein G magnetic beads (Active Motif, Cat. #53014). After stringent washes, immune complexes were eluted under non-denaturing conditions to preserve protein-DNA interactions. The sequences of primers used for ChIP-qPCR (human and mouse) in this study are listed in Supplementary Table S3 and S4.

### RNA-sequencing

MV4;11 cells were treated with vehicle DMSO (Control) or 108 nM ziftomenib or 165 nM selinexor or both for 24 hours. Cell pellets were submitted to LC Sciences on dry ice for poly(A)RNA-seq analysis (https://lcsciences.com/services). In brief, total RNA was extracted using a Qiagen RNA extraction kit (Qiagen; catalog no.1038703) following the manufacturer’s protocol. The quantity and purity were measured with Bioanalyzer 2100 and RNA 6000 Nano LabChip Kit (Agilent, CA, USA) with RIN number >7.0. Approximately 10 µg of total RNA was subjected to isolate Poly (A) mRNA with poly-T oligo-attached magnetic beads (Invitrogen). The poly(A)- or poly(A)+ RNA fractions are fragmented into small pieces using divalent cations under elevated temperature. Then, the cleaved RNA fragments were reverse-transcribed to create the final cDNA library (Illumina, San Diego, CA, USA), followed by paired-end sequencing on an Illumina Hiseq 4000. The raw data was cleaned by removing low-quality reads and adapters.

Genome mapping was performed using HISAT2 software followed by transcripts assembly by StringTie. All assemblies were merged to identify differentially expressed genes via mRNA expression profiling. StringTie^29^ was used to perform expression levels for mRNAs by calculating FPKM. The differentially expressed mRNAs were selected with log2 (fold change) >1 or log2 (fold change) <-1 and with statistical significance (p value < 0.05) by R package Ballgown^30^.

### Global proteomics

MV4;11 cells were harvested following treatment with 20 nM ziftomenib, 60 nM selinexor, and both for 24 hours and stored at −80 □. A total of 4 biological replicates for each treatment group (16 samples) were coded and submitted to the Wayne State University Proteomic Core for further processing. For global proteomic analysis, pellets from isolated cells were reduced/alkylated with DTT/IAA prior to MeOH precipitation using the Protifi protocol (ProtiFi, LLC; www.protifi.com). The precipitated proteins were washed with MeOH on S-trap cartridges, then resolubilized and digested with trypsin. Peptide abundance was determined using a fluorescent peptide quantitation kit (Pierce). Each digest was labeled with a unique TMT-Pro multiplexing reagent and then analyzed independently by LC-MS/MS on an Orbitrap Fusion or an Orbitrap QE MS system to ensure complete labeling. Once sample labeling was confirmed, the 16 samples were pooled together, and the mixture was fractionated by an alkaline reversed phase spin column (ThermoFisher, Waltham, MA, USA). Then, fractions were analyzed on the Orbitrap Eclipse using 2.0-hour cycle time analytical runs. All data were analyzed using Proteome Discoverer software. Additional analysis of the data set was performed using iPathwayGuide (version 2201; Advaita Bioinformatics, Plymouth, MI; http://advaitabio.com/ipathwayguide) to enable deeper interpretation of the differences between groups^31^.

### Cell line and patient-derived xenografts

NSG Mice used in these experiments were purchased from Jackson Laboratory (strain: 005557). The mice used in this study were cared for in accordance with a protocol approved by the Wayne State University Institutional Animal Care and Use Committee (IACUC-22-01-4305) and housed in sterile conditions.

In the first CDX model, 2×10^6^ GFP/luciferase-expressing MV4;11 cells were injected into each NSG mouse via the tail vein. On Day 8 after injection, mice were randomized to 4 cohorts after *in vivo* imaging confirmation of engraftment using the Bruker *in vivo* Xtreme imaging system. These mice (selinexor, n=4, ziftomenib, n=6, and Combo, n=6) were treated with ziftomenib (50mg/kg/5 days in a week) and selinexor (5mg/kg/biweekly) by oral gavage, and their combination for 21 days. All mice were weighed and monitored for the condition every day pre-and post-treatment and afterward for disease progression up to the end of the study. Mice were euthanized upon the manifestation of disease progression symptoms (persistent lethargy, anemia, eye lesion, internal mass involvement), and the survival curve was analyzed. In the second CDX experiment, a metronomic dosing regimen was employed to investigate how variations in dose intensity relate to the toxicity and efficacy of the combination therapy. The control group (n=4) was common to both the low and high-dose cohorts. The mice in the low- and high-dose cohorts were treated with ziftomenib (n=6, 50mg/kg/day and n=7, 100mg/kg for 5 days in a week) and selinexor (n=7, 2.5mg/kg/thrice weekly and n=8, 7.5mg/kg/biweekly) for 21 days and the combination respectively (n=7 and n=5). Mice without any drug were treated as controls in the whole experiment. The leukemic cells were analyzed for human (h) CD45^+^ cells in peripheral blood 2 weeks post-completion of treatment. For the OCI-AML3 CDX Model 1 (Survival Study): 3 × 10 GFP/luciferase-expressing OCI-AML3 cells were injected via the tail vein into each NSG mouse on Day 0. On Day 3 post-injection, mice were randomized into four treatment cohorts: control (n=7), selinexor (n=7), ziftomenib (n=6), and combination (n=7). Selinexor was administered by oral gavage at 5 mg/kg twice weekly (2 days apart), ziftomenib at 50 mg/kg/5 days in a week and the combination cohort received both agents on the same schedule. Control animals received vehicle on identical schedules. Treatment was continued for 28 days (Day 3-31). Humane endpoints included persistent lethargy, anemia, ocular lesions, or internal mass development. Upon reaching endpoint criteria, mice were euthanized, and survival data were used to construct Kaplan-Meier survival curves. In the second OCI-AML3 CDX experiment, to replicate an aggressive AML model with the intent to study the impact of treatment of leukemic load reduction, 2 × 10 GFP/luciferase-expressing OCI-AML3 cells were injected via the tail vein into each NSG mouse on Day 0. On Day 8, mice were randomized into four cohorts (n=5 per group): control, selinexor, ziftomenib, and combination. Treatment was initiated on Day 9. Selinexor was administered by oral gavage at 5 mg/kg twice weekly (Days 1 and 4 of each treatment week), ziftomenib at 50 mg/kg daily, and the combination cohort received both agents on the same schedule. Control animals received vehicle on identical schedules. Treatment continued for 14 consecutive days, including weekends. Representative bioluminescent imaging using Revvity IVIS^R^ imager was obtained to study the bioflux signals among the cohorts after the treatment period in the remaining surviving mice.

*KMT2A*-r PDX models were created using a patient-derived *KMT2A*-r primary AML cells (MLL-3 cells, Karmanos Cancer Institute), IV-engrafted into female immune-deficient NSG mice to evaluate the effectiveness of ziftomenib with selinexor, a therapeutic combination where both agents were administered orally with regimens of equivalent duration. 0.75 ×10^6^/mouse primary AML cells were injected into NSG mice via the tail vein. The experiment started with 40 mice in total, with 10 in each cohort. All mice were weighed and monitored for their condition every day pre- and post-injection and afterward for disease progression up to day 179. Mice were euthanized when progression symptoms presented (persistent lethargy, anemia, eye lesion, internal mass development). One mouse each from groups 1 and 3 was first euthanized from the study and terminally sampled for blood, spleen, and bone marrow on day 53 (confirmed to be progressive disease) after hind limb weakness. Three random mice from each group were removed from the study and terminally sampled blood, bone marrow, and spleen on the preplanned day 60 (harvested for further future analysis), leaving 7 mice for efficacy analysis as tabulated.

Ziftomenib was given orally daily at 50 mg/kg oral gavage starting day 5 through day 57 for a total dose of 135mg/kg. Mice were active, alert, and asymptomatic during the treatment phase. Selinexor was given PO at 10mg/kg/oral gavage on days 5, 8, and 11, and the dose was then de-escalated to 7.5mg/kg PO on day 14 with the Q3.5day schedule maintained for the remainder of treatment (concluding on day 56) for a total dose of 2,650mg/kg. Combination: ziftomenib and selinexor were given PO, matching their respective dose schedules as single agents above. Ziftomenib was given first, followed by selinexor with a 30-60 minutes split.

The *NPM1*-m AML7577 xenograft model was conducted at Crown Bioscience (San Diego, CA 92127). 2×10^6^ AML5777 (developed from the primary cells derived from a relapsed refractory *NPM1*-m AML patient) cells were injected into the tail vein of female NOD/SCID mice, and bone marrow was assessed weekly by FACS with human anti-CD45 antibody for tumor cell engraftment. Mice were randomized (day 0) into groups of 5 when bone marrow CD45+ count averaged 0.58%, and dosing was initiated 2 days later. Mice received ziftomenib at 25mg/kg PO QD, 50mgk/kg PO QD, or 100mg/kg PO/QD or selinexor at 5mg/kg PO BIW, or the combination of one of the ziftomenib doses with selinexor for 30 days. The mice continued to be observed until day 70, at which point the study was terminated on 70 days and bone marrow was collected for hCD45% assay by FACS (hCD45 and L-D), spleens were weighed, and livers and lungs were collected for FFPE blocks and used for CD45 IHC analysis.

### Flow cytometric detection of human cells in mouse cheek bleeds

Heparinized blood from cheek bleeds (100-200uL) was collected, and mononuclear cells were obtained by brief whole blood lysis in 3mL sterile distilled water followed by washing in RPMI-1640 medium supplemented with 10% FBS which contained 0.3 mL 10X PBS to restore tonicity. An additional round of lysis was performed if evidence of red cells was observed in the pelleted cells. Cells were resuspended in a complete RPMI medium and stained with anti-human CD45-PC5 (Beckman Coulter; Cat# IM2653U) to assess human cell presence. Additional human specific monoclonal antibodies were added as necessary to further characterize human cell subset populations. The percentage of human cells was quantified by several specific human-related CD marker positivity events in relation to total mononuclear events obtained. Resident mouse lymphocytes served as an internal negative control. Flow cytometry was performed on a Beckman Coulter XL-MCL unit equipped with a 488nm Argon laser, and events were analyzed by System II Software.

### Immunohistochemical analysis

Post-necropsy immunohistochemical analysis (IHC) of mice liver and lung tissues was performed for human CD45 (Rabbit anti-human, Cell Signaling Technology, Catalog no. 13917S). The secondary antibody detection kit was Goat anti-Rb IgG (Leica, Catalog no. DS9800). Prior to staining, all the tissues from mice were stored in 10% neutral-buffered formalin (v/v). IHC was carried out following standard techniques. Paraffin sections were de-waxed and rehydrated in a xylene-ethanol series. Endogenous peroxides were removed by a 3-4% (v/v) hydrogen peroxide solution and incubated at room temperature for 25 minutes. Antigen retrieval was done with a pH 9.0 EDTA buffer at 100 for 20 minutes. The primary antibody dilutions were 1:800. A secondary antibody detection kit was used for detection (Leica, Catalog no. DS9800). All the images were analyzed with the HALOTM platform at 20x magnifications.

The whole slide image was analyzed, and considerable necrosis was excluded. Both the total tissue area and the number of IHC-positive cells were counted. IHC scores were presented as the ratio of the positive cell counts against the total tissue area. IHC-positive cell density = Positive cell counts / Total tissue area.

### Statistical analysis

The experiments were done at least in triplicates. Data represented as mean ± standard deviation (SD). The data were compared using Student’s t-test. The *p* value of < 0.05 was considered as statistically significant. Symbol * indicates *p* < 0.05, ** indicates *p* < 0.01 and *** indicates *p* < 0.001.

## Results

### 3.1 Ziftomenib and selinexor inhibit the growth of *KMT2A*-r and *NPM1*-m AML cells and enhance cell death

An analysis using a network topology computational system approach (http://slorth.biochem.sussex.ac.uk) revealed that MEN1 and XPO1 interact via multiple signaling molecules and targeting them together could be synthetically lethal. The lethality score was high at 0.88 out of 1 (Figure 1A and Supplemental Figure S1). In this study, the menin inhibitor ziftomenib and XPO1 inhibitor selinexor and their combination were tested *in vitro* in multiple *KMT2A*-r AML cell lines (MV4;11, MOLM13, and SEM) and two *NPM1*-m AML cell lines (OCI-AML3 and IMS-M2). We observed noticeable growth inhibition with ziftomenib and selinexor in these cell lines (Figure 1B, Supplemental Figure S2A-D). Synergy analysis revealed the combination of ziftomenib and selinexor on these cells was synergistic in a wide range of doses (Figure 1C-F). Most combination index (CI) values were below 1 and within the synergy triangle of isobolograms (Figure 1G&H and Supplemental Figure S2E-J). We observed similar synergistic trends in the SEM cell line (Supplemental Figure S2K-O). To evaluate the mode of cell death, flow cytometric analysis of annexin-V/PI-stained cells revealed an increase in early apoptotic cell deaths in *KMT2A*-r AML cell lines in ziftomenib (10.3%) and selinexor (4.5%) groups compared to control (2.1%) (Figure 1I). Similarly, apoptosis was observed in *NPM1*-m AML cell lines in ziftomenib (7.3%) and selinexor (27.8%) groups compared to the control (3.9%) (Figure 1J). When the inhibitors were combined, early apoptosis increased to 14.7% in *KMT2A*-r and 45% in *NPM1*-m AML. The significant increase in apoptosis observed with the combination of ziftomenib and selinexor suggests a synergistic interaction between these two compounds. Compared to controls, ziftomenib- and selinexor-treated samples showed an increase in the G0/G1 population and a decrease in S, G2/M populations, which was significantly enhanced in both the molecular signatures when the treatments were combined (Supplemental Figure S2P&Q), suggesting a robust modulation of the cell cycle proteins by this combination.

**Figure 1.**
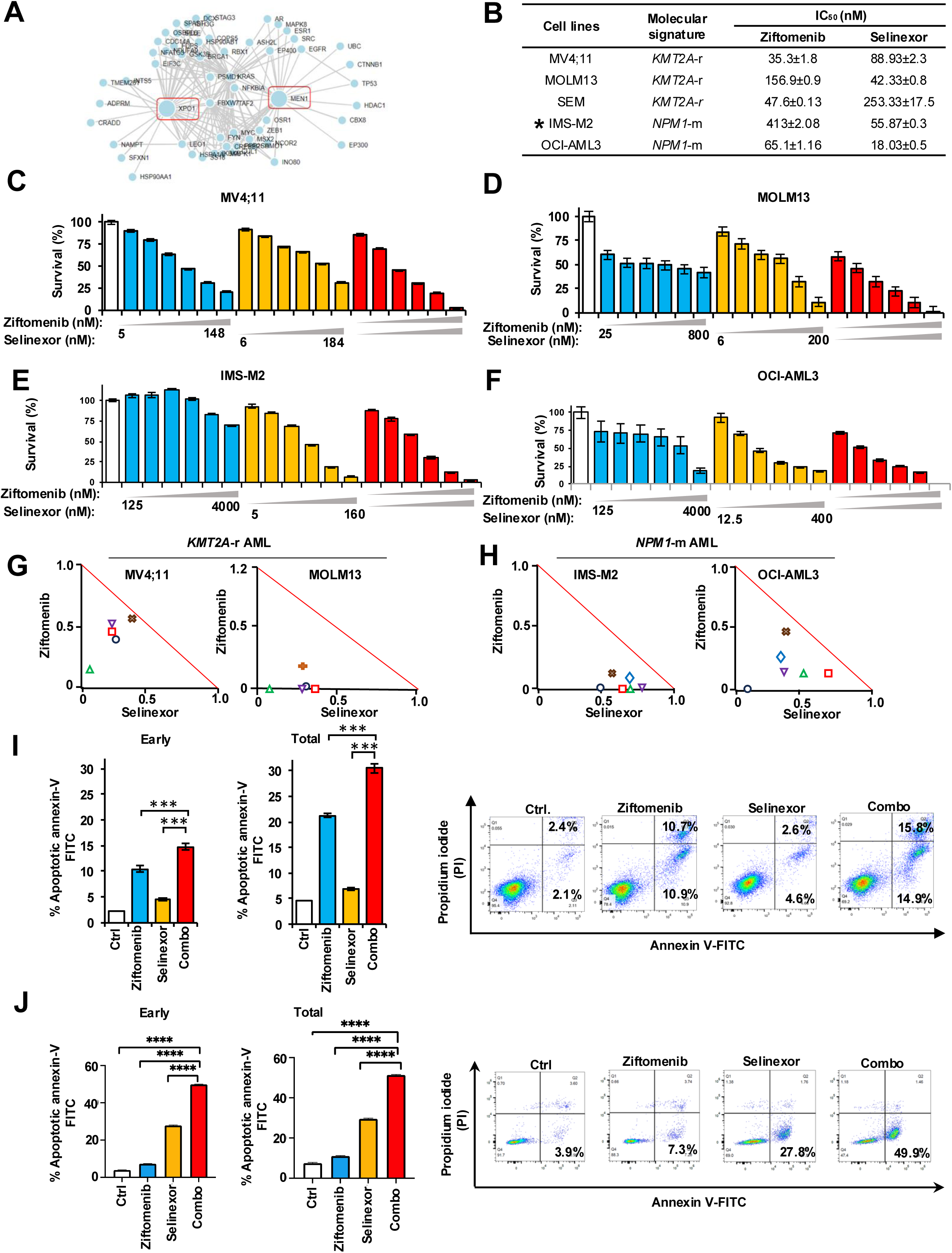
*In vitro* activity of ziftomenib and selinexor on *KMT2A*-r and *NPM1*-m AML cell lines. (A) Network topology map generated with Slorth (www.slorth.biochem.sussex) illustrating direct and indirect molecular interactions between MEN1 and XPO1. (B) IC_50_ values of ziftomenib and selinexor in different *KMT2A*-r and *NPM1*-m AML cell lines. (C-F) Ziftomenib and selinexor combination treatment in MV4;11 and MOLM13 *KMT2A*-r and IMS-M2 and OCI-AML3 *NPM1*-m AML cell lines. (G-H) Normalized isobolograms for MV4;11 and MOLM13 *KMT2A*-r and IMS-M2 and OCI-AML3 *NPM1*-m AML cell lines. Isobolograms were generated using Calcusyn 2.1 software. (I-J) Apoptotic cell deaths in MV4;11 and OCI-AML3 cells after ziftomenib, selinexor and combination treatments. Apoptotic cells were determined using annexin V-propidium iodide (PI) flow cytometric assay. DMSO or drug treatments were performed for 48-72 hours. Representative flow cytometric images are shown in the right panel. ***, *p* < 0.001.

### 3.2 Ziftomenib and selinexor combination is active in primary *KMT2A*-r and *NPM1*-m AML cells

The combination of menin and XPO1 inhibitors measurably suppressed the survival of patient-derived *KMT2A*-r and *NPM1*-m primary CD34+ progenitor stem cells and their ability to form colonies in *KMT2A*-r primary cells. We observed robust synergy (CI<1) in several *KMT2A*-r (n=9) and *NPM1*-m (n=5) primary AML cells in culture (Figure 2A and B) and across multiple dose ranges (CI<1, suggesting synergy, Supplemental Table S5 and S6). More than 50% of primary *KMT2A*-r AML cells died in the combination group (Figure 2C). The cytogenetics, translocations and mutations of the KMT2A-r/*NPM1*-m primary leukemic samples are shown in Supplemental Table S7 and S8. The MethoCult assay demonstrated a substantial reduction in the number of colonies derived from human CD34 positive primary patient derived *KMT2A*-r cells in the combination group at 14 days, indicating a robust inhibitory effect of the combined treatment on colony formation (Figure 2D). Colony morphology from human CD34 positive primary patient derived *KMT2A*-r cells was also disrupted by combined menin and XPO1 inhibitors in high and low doses (Figure 2E and F). Morphologically, colony phenotypes in the combination group were noticeably less dense than single-agent-treated cells (Supplemental Figure S3A and B). Importantly, under the same treatment conditions, no significant toxicity was observed in normal CD34+ human hematopoietic progenitor stem cells (HPSCs) obtained from healthy donors even after 72 hours of culture (Figure 2G). These findings indicate that combining menin and XPO1 inhibitors selectively targets these molecular signatures synergistically, warranting further investigation.

**Figure 2.**
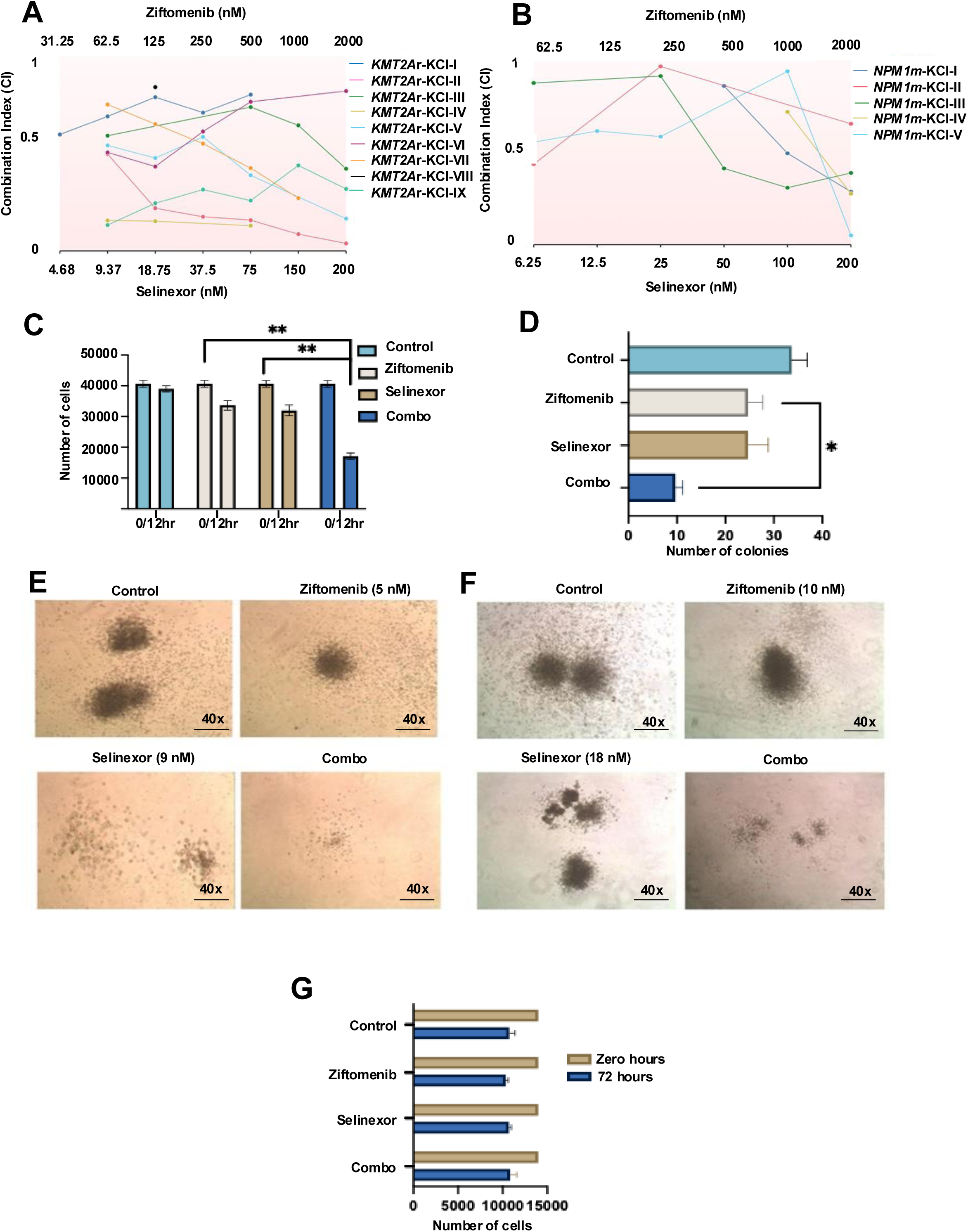
Ziftomenib and selinexor combination is active in primary *KMT2A*-r and *NPM1*-m AML cells. (A-B) Consolidated combination index of multiple patient-derived primary CD34+ *KMT2A*-r and primary *NPM1*-m leukemic cells treated with different doses of ziftomenib and selinexor. (C) Inhibition of CD34+ primary *KMT2A-*r AML cells growth after drug treatment with the doses indicated for 12 hours. The y-axis indicates the number of cells. (D) Inhibition of colony formation read post-14 days in culture after menin inhibitor and selinexor treatments with the doses indicated. The x-axis indicates the number of colonies. (E&F) Representative images of the colonies after 14 days in culture of primary *KMT2A*-r primary cells with treatment. (G). Effect of menin inhibitor and selinexor on the normal human CD34^+^ stem cells. The x-axis indicates the number of cells, and y-axis indicates the treatments. *, *p* < 0.05.

### 3.3 Menin and XPO1 combinatorial inhibition impacts menin-controlled gene expression programs

The quantitative PCR data revealed the downregulation of several menin targets, including *HOXA10*, *MEIS1*, and *PBX3*, in the combination treatment group in both *KMT2A*-r and *NPM1*-m models (Figure 3A and B respectively). Western blot analysis showed that ziftomenib and selinexor, either alone or in combination, downregulated menin and its target gene expression, including HOXA9, HOXA10, and MEIS1, consistent with the qPCR findings (Figure 3C&D and Supplemental Figure S4A-J) in both models. In addition, the combination increased CD11b expression, a marker of monocytic differentiation, in treated MV4;11 and OCI-AML3 cells (Figure 3C&D and Supplemental Figure S4E&J). Ziftomenib and selinexor suppressed menin expression in a time-dependent manner, with reduction evident as early as 3 hours of treatment which is more pronounced in the combined (Supplementary Figure S5A). MV4;11 cells were treated with ziftomenib, selinexor, and their combination for 24 hours followed by a drug washout period. The downregulation of HOXA10 and MEIS1/2 from the drug combination was more sustained at zero, 6-, and 18-hours post-washout (Supplementary Figure S5B and S5C). To further characterize the transcriptional impact, MV4;11 cells were treated with ziftomenib, selinexor, or their combination, and RNAseq and proteomic analyses were performed. An unbiased approach revealed more genes to be differentially expressed in the combination than any single agent (Figure 3E and Supplemental Figure S6A&B). The gene ontology (GO) enrichment revealed several terms, including cell cycle, cell division, DNA repair, and DNA replication (Figure 3F). Gene set enrichment analysis (GSEA) demonstrated a profound reduction in the enrichment score of the cell cycle following combination treatment (Supplemental Figure S6C). A curated set of genes significantly altered by ziftomenib treatment, including canonical menin/KMT2A targets, was more downregulated in the combination treatment (Figures 3G). These transcriptional changes aligned with previously defined MLL/menin-bound loci^32^ (Figure 3H). Notably, key oncogenic drivers such as *HOXA* cluster genes and *MEIS1* were downregulated, consistent with impaired menin/KMT2A activity.

**Figure 3.**
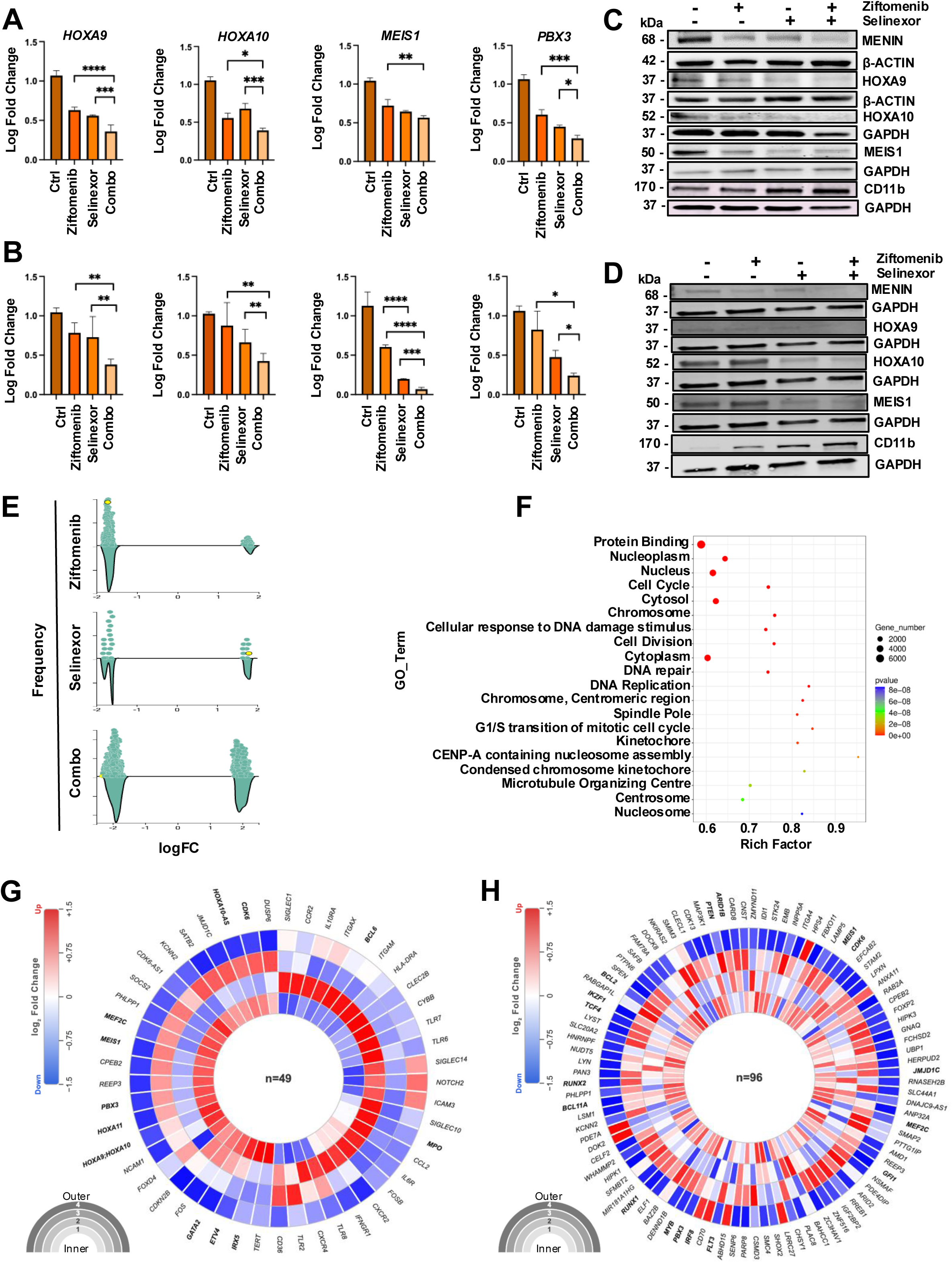
Menin and XPO1 combinatorial inhibition impacts menin-controlled gene expression programs. (A&B) Quantitative expression of menin downstream targets in MV4;11 and OCI-AML3 cell lines as determined by real-time qPCR after 24 hours of treatment with ziftomenib (50 nM), selinexor (100 nM), or combination. (C&D) Expression of menin and its downstream target proteins, including HOXA9, MEIS1, when treated for 72 hours with either DMSO or 36 nM ziftomenib and 55 nM selinexor in MV4;11 and OCI-AML3 cells. (E) Global expression of mRNAs in the indicated treatment groups. (F) Affected gene ontology (GO) terms from GO analysis. (G) Circular heatmap of curated menin/KMT2A target gene expression across treatment conditions displays a curated set of bona fide menin/KMT2A/menin inhibitor-responsive genes identified from RNA sequencing. Expression patterns of these genes are shown across control, ziftomenib monotherapy, selinexor monotherapy, and ziftomenib-selinexor combination treatment groups in MV4;11 cells. Both upregulated and downregulated transcripts are highlighted, demonstrating the selective transcriptional modulation of menin/KMT2A target programs and the enhanced impact of combination therapy. (H) Differential expression of biologically relevant genes defined by heavy MLL/menin-bound loci^32^ and impact of different treatments on those genes in MV4;11 treated cells. Concentric rings 1 to 4 represent expression changes across four conditions (inner to outer: Control, Ziftomenib, Selinexor, and Combination), with color intensity reflecting log fold change (range −1.5 to +1.5).

Importantly, the combination of ziftomenib and selinexor produced a more pronounced disruption of these transcriptional programs, supporting their mechanistic synergy. Each circular heatmap displays differentially expressed genes (Figure 3G, n=49; Figure 3H, n=96), with genes in bold italic denoting loci with established roles in HOX/MEIS-driven transcriptional regulation and menin/KMT2A complex activity.

Proteomic analysis also identified cell cycle signaling pathways to be perturbed after ziftomenib and selinexor treatment (Supplemental Figure S6D&E). All the differentially expressed proteins in the cell cycle, including CCNB2, TTK, CDKN2A, and CDK4, were found to be downregulated.

### 3.4 XPO1 regulates menin-KMT2A complex formation, chromatin occupancy and transcriptional function

Menin is known to contain two NES motifs capable of binding XPO1^33^, however, selinexor treatment did not increase nuclear accumulation of menin (Supplemental Figure S7A), suggesting that the interaction may not be important for the nuclear exportation of menin in myeloid progenitor cells. Recent studies have suggested that XPO1 can function as a transcriptional regulator^22, 24, 34^. We therefore investigated the possibility that this interaction may be critical for the transcriptional activity of menin in cells transformed by KMT2A fusions. To exclude possible interference from unknown mutations in the human leukemia cell lines, we first examined this possibility in mouse myeloid progenitors freshly immortalized by transduction of *KMT2A*/*MLLT3*-expressing retrovirus. To test whether blocking its interaction with XPO1 may affect the ability of menin to bind to chromatin, we treated these cells with KPT-185, which has higher potency than selinexor *in vitro*^35^, for a short period of 6 hours, and subsequently examined menin occupancy at the *Hoxa9* locus by chromatin immunoprecipitation (ChIP) analysis (Figure 4A). Interestingly, a significant reduction in menin-binding was detected at different regions of *Hoxa9* after KPT-185 treatment, while no significant decrease in menin protein levels was observed in the nuclear extracts of treated cells (Supplemental Figure S7A), suggesting that the interaction between menin and XPO1 may be critical for the binding of menin to chromatin.

**Figure 4.**
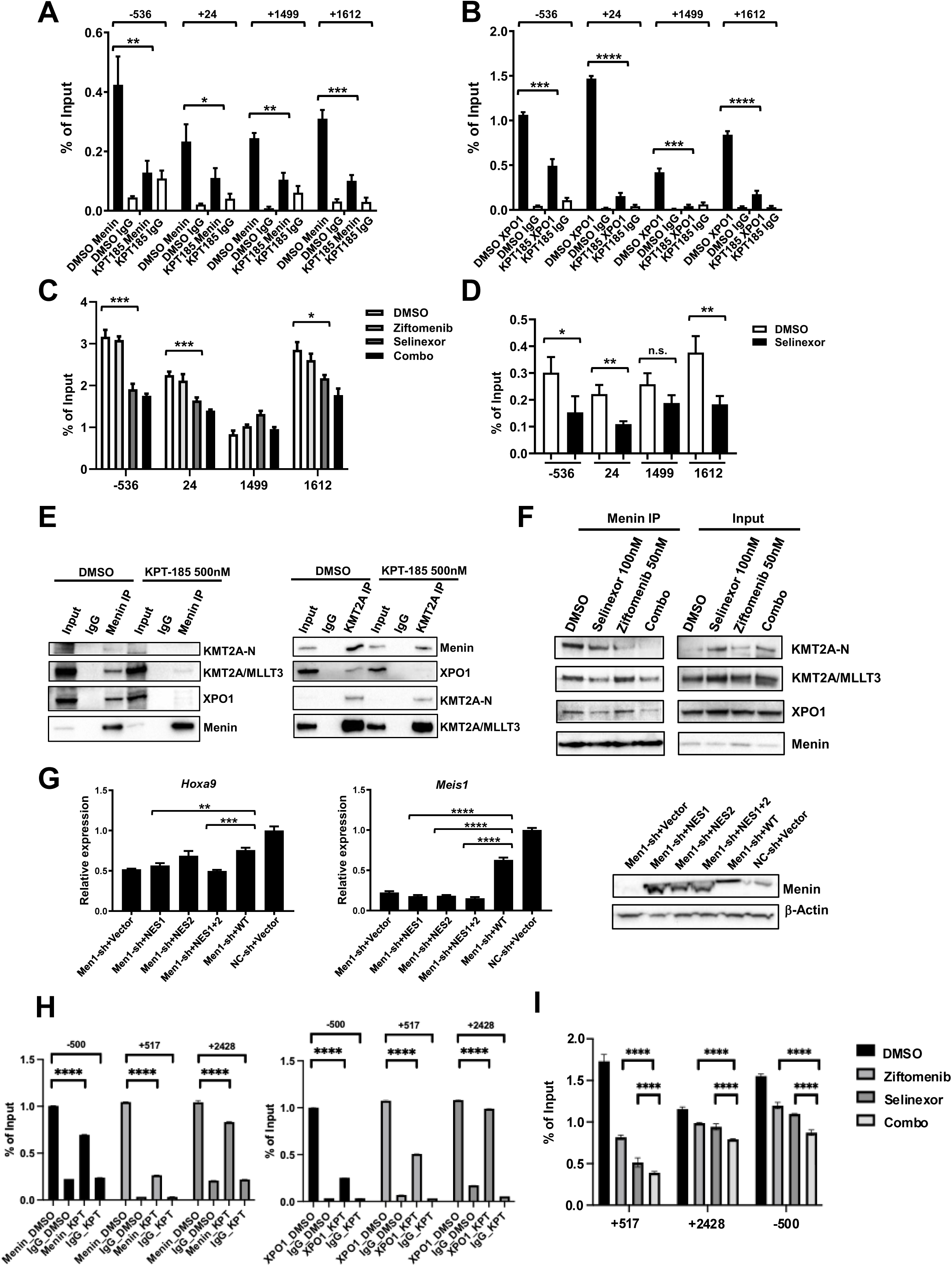
XPO1 is critical for stable binding of menin to chromatin and KMT2A/KMT2A fusions. (A) Representative ChIP-qPCR analyses of indicated regions of *Hoxa9* locus relative to the transcriptional start site (TSS) using an anti-menin antibody or IgG in KMT2A/MLLT3-immortalized myeloid progenitor cells treated with KPT-185 (500 nM) or DMSO for 6 hours. (B) Representative ChIP-qPCR analyses of indicated regions of *Hoxa9* locus using an anti-XPO1 antibody in same cells as in (A). (C) Representative ChIP-qPCR analyses of indicated regions of *Hoxa9* locus using an anti-menin antibody or IgG in KMT2A/MLLT3-immortalized myeloid progenitor cells treated with ziftomenib (50 nM) alone, selinexor (100 nM) alone, the combination, or DMSO for 6 hours. Percent of Input values for menin are presented after deduction of values for IgG. (D) Representative ChIP-qPCR analyses of indicated regions of *Hoxa9* locus using an anti-XPO1 antibody in same cells as in (C) treated with selinexor (100 nM) or DMSO for 6 hours. Percent of Input values for XPO1 are presented after deduction of values for IgG. (E) Left panel, immunoprecipitates were prepared from nuclear extract of HEK293T cells transiently transfected with pMSCVpuro-KMT2A/MLLT3 using a menin-specific antibody or control IgG and analyzed by Western blotting analysis using indicated antibodies. Right panel, immunoprecipitates prepared from same nuclear extract using a KMT2A-N-specific antibody or IgG were analyzed by Western blotting analysis. (F) Immunoprecipitates were prepared from nuclear extract of HEK293T cells transiently transfected with pMSCVpuro-KMT2A/MLLT3 and treated with ziftomenib (50 nM) alone, selinexor (100 nM) alone, the combination, or DMSO for 6 hours using a menin-specific antibody and analyzed by Western blotting analysis using indicated antibodies. (G) Real-time RT-PCR analysis of *Hoxa9* (left panel) and *Meis1* (right panel) mRNA levels in KMT2A/MLLT3-immortalized myeloid progenitors transduced with indicated empty pMYs retrovirus (Vector) or pMYs virus expressing menin NES mutants (NES1, NES2, and NES1+2) in combination with a lentiviral shRNA targeting 3’UTR of endogenous *Men1* (Men1-sh) or a non-targeting shRNA (NC-sh). Bottom panel, Western blotting analysis of menin expression in the co-transduced cells. (H) MV4;11 cells were treated with selinexor (or vehicle control, as indicated), followed by ChIP-qPCR using an anti-menin antibody (left panel) and anti-XPO1 antibody (right panel). qPCR was performed with primers spanning the indicated regions across the *HOXA9* locus. ChIP enrichment is plotted as percent input (% input) for each amplicon; IgG served as a negative control. Bars represent the measured enrichment for each condition. (I) ChIP-qPCR using an anti-menin antibody in MV4;11 cells to map occupancy across the HOXA9 locus following treatment with vehicle control, ziftomenib, selinexor, or the combination.

Consistent with the reduction in menin-binding to chromatin after KPT-185 treatment, the mRNA levels of *Hoxa9*, *Hoxa10*, *Meis1* were also significantly reduced at this time point (Supplemental Figure S7B). It was shown previously that treatment with selinexor at 1 µM concentration for 24 hours can also reduce the binding of XPO1 to chromatin^34^. We therefore examined XPO1-binding to the *Hoxa9* locus in the same cells by ChIP. A significant reduction in XPO1-binding was also observed after KPT-185 treatment, further suggesting that XPO1 may be required for the binding of menin to chromatin (Figure 4B). Remarkably, selinexor at a much lower concentration of 100 nM also significantly decreased the binding of menin at *Hoxa9* in these cells without affecting its levels in the nucleus (Figure 4C and Supplemental Figure S7C), which, as expected, was further reduced in cells treated with the combination of selinexor and ziftomenib (Figure 4C). The occupancy of XPO1 at *Hoxa9* was also reduced by selinexor at this concentration (Figure 4D). Consistent with these results, the mRNA levels of *Hoxa9*, *Hoxa10*, *Meis1, and Pbx3* were also reduced (Supplemental Figure S7D).

We also examined the binding of KMT2A and KMT2A/AF9 at the same sites after KPT-185 treatment by ChIP using a KMT2A-N-specific antibody. Interestingly, there was no reduction detected at the same sites, further suggesting that the complex formation between menin and KMT2A may be disrupted due to the loss of XPO1-binding to menin (Supplemental Figure S7E). To investigate this notion further, we treated HEK293T cells transiently expressing KMT2A/MLLT3 similarly with KPT-185 or DMSO for 6 hours and performed immunoprecipitation of menin from nuclear extracts of these cells. As expected, both MLL/AF9 and XPO1 were pulled down together with menin in DMSO-treated cells. In contrast, significantly less of both proteins were detected in the menin immunoprecipitates from KPT-185-treated cells, despite more menin protein being pulled down (Figure 4E, left panel). Significantly less menin was also pulled down by a KMT2A-N-specific antibody after KPT-185 treatment (Figure 4E, right panel). The amounts of KMT2A-N and KMT2A/MLLT3 interacting with menin were also reduced by treatment with 100nM selinexor and were further reduced by the additional treatment with ziftomenib (Figure 4F). These results further support the idea that the interaction between XPO1 and menin may be critical for stabilizing the interaction between menin and wild-type KMT2A and KMT2A/MLLT3.

To exclude potential interference from possible secondary effects induced by blocking XPO1-mediated nuclear exportation, we generated menin mutants with either one or both NES motifs mutated (Supplemental Figure S7F). Consistent with the minimal effects of XPO1 inhibitors on the nuclear levels of menin, these mutant proteins did not display any significant changes in their nuclear/cytoplasmic localization compared to the wild-type protein when they were transiently expressed in HEK293T cells (Supplemental Figure S7G). However, only the NES2 mutant caused an increase in Hoxa9 mRNA levels and none of the mutants was able to upregulate *Meis1* expression in *KMT2A/MLLT3*-immortalized cells after the endogenous *Men1* was knocked down using a shRNA targeting its 3’UTR (Figure 4G). Consistent with these results, all three mutants failed to increase colony formation after the *Men1* knockdown (Supplemental Figure S7H). These results further support the idea that the interaction with XPO1 is critical for menin to activate target gene transcription.

To test whether the interaction with XPO1 is critical for chromatin recruitment of menin in human AML cells, we also performed ChIP analyses in MV4;11 cells after treatment with selinexor for 12 hours. Similar to the results from *KMT2A/MLLT3*-immortalized mouse myeloid progenitors, significant reductions in menin and XPO1 occupancy were also detected at *HOXA9* after selinexor treatment (Figure 4H, Supplemental Figure S7I) in MV4;11 cells and menin occupancy was further reduced after additional treatment with ziftomenib (Figure 4I).

In combination, these results strongly suggest that the interaction with XPO1 is critical for menin to form stable complexes with KMT2A and KMT2A/MLLT3 and to bind chromatin. More efficient disruption of these complexes and their binding to chromatin by the combination of ziftomenib and selinexor may significantly contribute to their synergy in inhibiting *KMT2A*-r and *NPM1*-m induced leukemia cells.

### 3.5 Ziftomenib and selinexor combination shows anti-leukemia activity in cell line- and patient-derived xenograft (CDX/PDX) models *in vivo*

MV4;11 cell line-derived xenografts in NSG mice showed a significant difference in survival in the combination arm compared to any single agent, with no overt toxicity as evidenced by stable weights throughout the treatment period (Figure 5A and Supplementary Figure S8A). These findings were biologically validated in a metronomic dosing experiment using varying higher and lower doses of selinexor and ziftomenib to assess the impact of dose variations on efficacy and toxicity, which similarly demonstrated a robust survival advantage in the combination cohort (Figure 5B, 5C, Supplementary Figures S8B and S8C). Circulating human (h) CD45 positive cells as measured by flowcytometry (FCM) 2 weeks post completion of treatment in all groups, showed deeper response in the combination-treated mice (Figures 5D, 5E and 5F). In NSG mice bearing *NPM1*-m (OCI-AML3) xenografts, combination therapy resulted in significantly improved survival relative to single-agent treatment (Figure 5G). With the intent of measuring the impact of treatment leukemic load reduction, the combination indeed showed a significant reduction in the total BioFlux compared to any single agents in NSG mice with GFP/Luc+ OCI-AML3 (*NPM1*-m) cell line-derived xenograft mice (Figure 5H and Supplemental Figure S8E).

**Figure 5.**
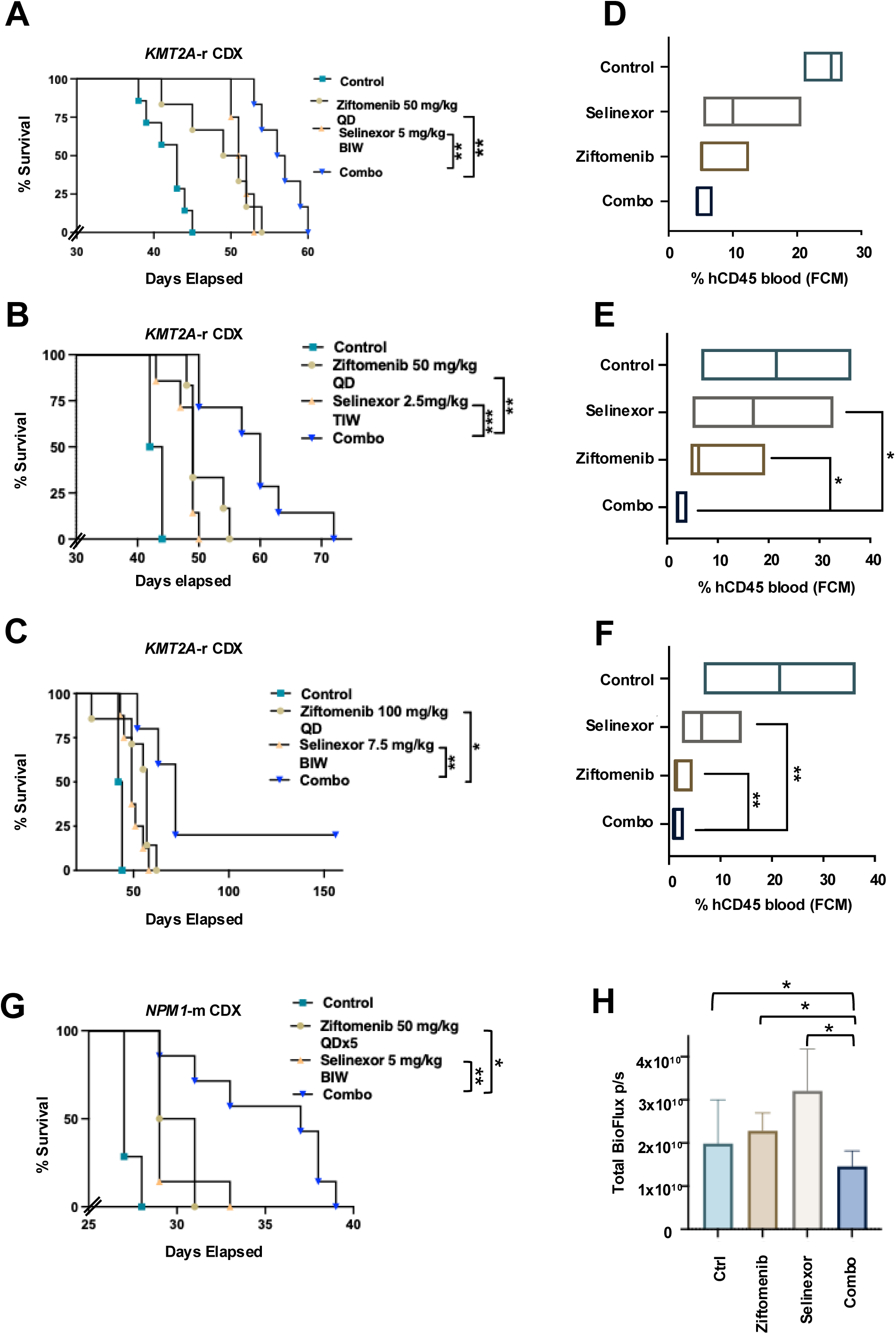
Effect of menin inhibitor and selinexor combination in MV4;11 and OCI-AML3 cell line-derived xenograft (CDX) models. (A-C) Survival of MV4;11 cell line-derived xenograft mice treated with metronomic doses of the ziftomenib and selinexor, and in combination. (D-F) Percentage of human CD45 positive cells in the peripheral blood of MV4;11 cell line-derived xenograft mice groups two weeks post last treatment (3 mice/group) with different doses of ziftomenib and selinexor. (G) Survival of GFP/Luciferase positive OCI-AML3 cell line-derived xenograft mice treated with ziftomenib and selinexor, and with combination. (H) Bioluminescence from luciferase in different groups of GFP/Luciferase-positive OCI-AML3 cell line-derived xenograft mice.

Patient-derived xenograft NSG mice model using human primary *KMT2A*-r retransplantable leukemia cells showed superior survival in the combination arm without signs of overt toxicity, as reflected by stable body weights during the treatment course (Figure 6A and Supplemental Figure S8D). Three of 7 mice in the ziftomenib group had no evidence of disease progression on the last day of the study (day 179), yielding a 113% increase in lifespan (%ILS) for this cohort. No long-term symptom-free mice emerged from only selinexor treatment, which produced a modest 11% ILS. The combined treatment produced five out of seven mice that remained symptom-free at the end of the study period, resulting in an increase of over 129% ILS. The exact percentage of ILS could not be determined due to the inability to calculate this group’s median day of death. These findings suggest, at the very least, a signal to synergy for that PDX model, though it may not be statistically significant because of the sample numbers and the study design. A more robust group size is needed to determine if drug interaction is synergistic in this model. Patient-derived xenograft models using human primary *NPM1*-m AML similarly showed significant synergy in the combination cohort (Figure 6B and Supplemental Figure S8F), with an early and marked reduction in the circulating hCD45 with the combination treatment, an effect that was sustained in this group even after 2 weeks of treatment discontinuation (Figure 6C).

**Figure 6.**
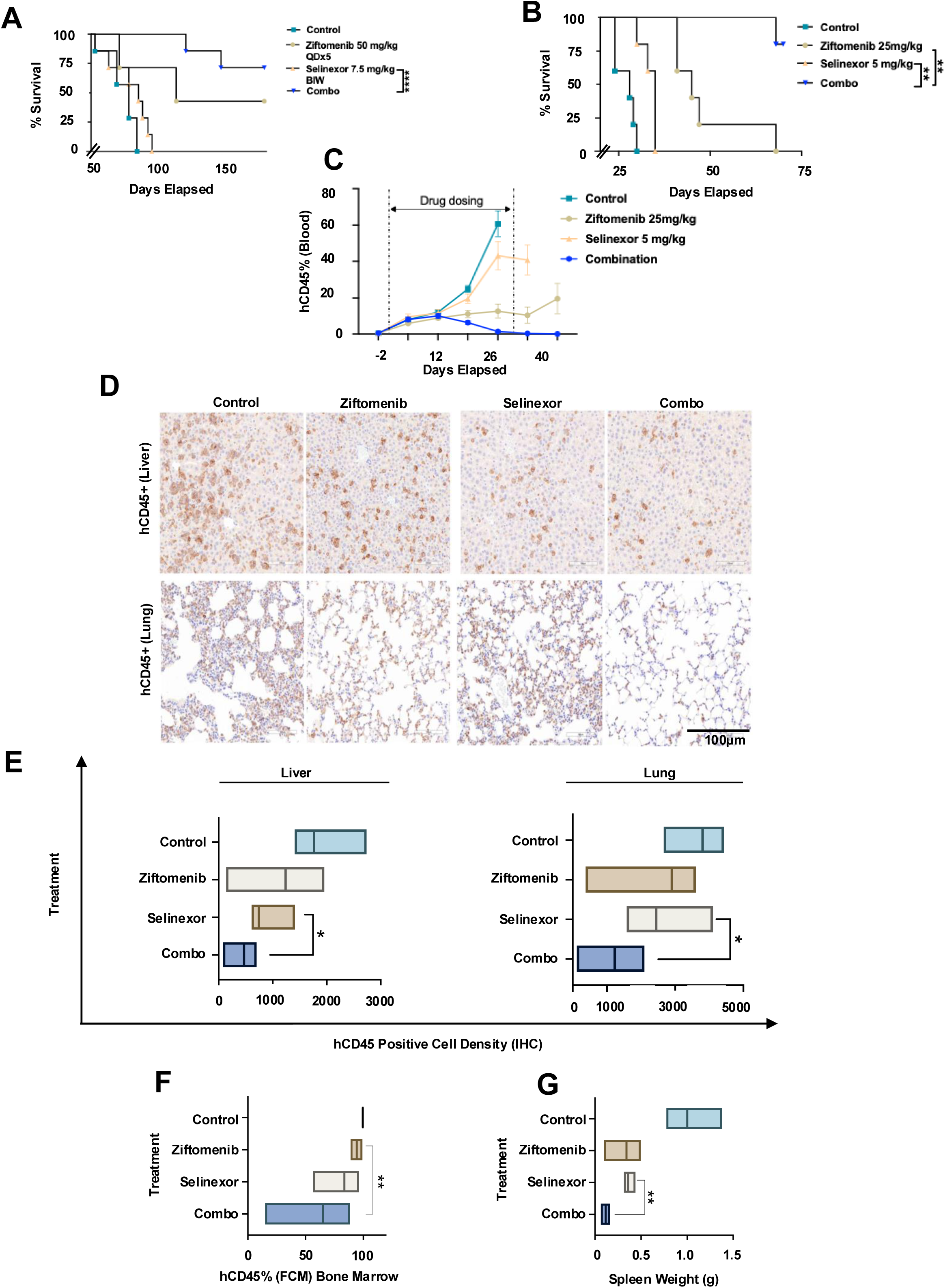
Effect of menin inhibitor and selinexor combination in patient-derived xenograft (PDX) models. (A-B) Survival of vehicle or inhibitor-treated *KMT2A-*r and *NPM1*-m PDX mice. (C) Residual leukemia evaluation on treatment and post-necropsy in *NPM1*-m PDX NSG mice. (D) Human (h) CD45 staining in liver biopsy (top panel) and lung biopsy (bottom panel) under the vehicle control, ziftomenib, selinexor, and combination treatment cohorts. (E) hCD45+ cell density score (IHC) in Liver and Lung tissues. (F) Residual Bone marrow hCD45% by FCM during and after treatment in control, ziftomenib, selinexor, and combination cohorts. FCM, flow cytometry. (G) Measured spleen weight post-necropsy/termination in NSG mice in all cohorts

Post-necropsy hCD45 cell densities measured by immunohistochemistry in mice liver and lung were consistently and significantly recorded as low in the combination treatment cohort, suggesting a robust control of extramedullary infiltration of leukemic cells (Figures 6D&E). hCD45 measured in the bone marrow and the spleen weight in the combination-treated mice were significantly less compared to the single agent-treated group (Figures 6F&G).

## Discussion

Menin inhibitors are emerging as promising therapies in *KMT2A*-r and *NPM1*-m AML. Biomarkers such as HOX or MEIS1 expression could help identify additional AML subsets that can be sensitive to MI. However, recent studies using revumenib have identified MEN1 binding-site ‘gatekeeper’ mutations and other noncanonical genetic and epigenetic processes that mediate resistance to MI^14, 36^. These findings emphasize the need for rational combination strategies to enhance the depth and durability of response while mitigating resistance.

Here, we demonstrate that the combined inhibition of menin and XPO1 with ziftomenib and selinexor is synergistically anti-leukemic across multiple *KMT2A*-r and *NPM1*-m AML models. Dual inhibition suppressed AML cell growth, diminished colony-forming capability of CD34+ leukemic progenitors, increased differentiation and markedly increased apoptosis while sparing normal hematopoietic CD34+ stem cells. *In vivo*, combined targeting showed significantly increased survival advantages in multiple CDX and PDX models, including under metronomic low-intensity dosing, underscoring its translational potential.

Mechanistically, our work identifies a critical functional dependency between menin, KMT2A/KMT2A-fusion complexes, and XPO1-associated transcriptional machinery. Interaction with XPO1 appears critical for menin to bind to chromatin, as XPO1 inhibition significantly reduced menin’s chromatin occupancy. This parallels the previously known effect seen with MIs, suggesting that menin’s recruitment to chromatin requires its interaction with both KMT2A and XPO1. Additionally, XPO1 interaction is also crucial for stabilizing menin-KMT2A and menin-KMT2A/MLLT3 complexes as XPO1 inhibition quickly disrupted these complexes as shown in our co-IP studies. Although these observations may reflect independent events, reduced menin chromatin occupancy could also be from the disruption of these complexes. XPO1 interaction has been established as a critical requirement for aberrant transcriptional activity of several mutant proteins implicated in leukemogenesis^23–25, 37^. Our finding identifies wild-type menin, similar to wild-type SETBP1^24^, also depends on XPO1 for its transcriptional regulatory function, suggesting that XPO1 contributes to normal transcriptional regulation in addition to its canonical role as a nuclear exporter.

Our results also suggest that XPO1 engages with chromatin through its NES-binding cleft. We were able to detect a significant reduction in XPO1 binding to chromatin after a much shorter exposure with selinexor (6 hours) at a much lower concentration (100 nM) than what was previously reported^25^, suggesting a direct chromatin association of XPO1 mediated by its NES-binding cleft than the consequence of the nuclear export inhibition per se. Although the molecule that recruits XPO1 to chromatin remains to be identified, it is possible that the differences in the effective selinexor concentrations to block XPO1-binding to chromatin between ours and other studies may reflect the variable accessibility of the inhibitor to its target site, governed by the variations in chromatin-bound factor among different cell types. Given that XPO1 can dimerize when exporting cargos^38^, it is conceivable that menin’s chromatin binding may require a dimeric form of XPO1.

The observation that menin’s interaction with XPO1 is critical for its chromatin binding and for its stable complex formation with KMT2A and KMT2A/MLLT3 provides a mechanistic basis for the synergy between ziftomenib and selinexor in inhibiting *KMT2A*-r leukemic cells.

Targeting the KMT2A and KMT2A-fusion complexes through distinct mechanisms with the combination possibly more efficiently disrupts the complexes and their chromatin association, ultimately leading to a greater reduction in their target gene transcription. Moreover, simultaneous disruption of two discrete protein-protein interaction interfaces within the KMT2A/KMT2A-fusion assemblies is anticipated to deepen response and delay relapse that may likely develop through the menin inhibitor binding site mutations or other non-canonical pathways.

Synergistic growth inhibition was consistent observed in several *KMT2A*-r and *NPM1*-m AML models, suggesting that menin and XPO1-associated pathways converge on shared transcriptional programs. Further, our preliminary work has elucidated the role of XPO1 in menin occupancy at multiple HOX clusters in the chromatin, suggesting a critical reason for synergy in *KMT2A*-r leukemia. The combination robustly induced apoptosis and differentiation, including upregulation of CD11b, consistent with the enhanced suppression of HOX/MEIS1-driven leukemogenic program. These findings align with previous reports showing that menin inhibition promotes a tumor-suppressive transcriptional reprogramming^28^ and suggest that XPO1 inhibition amplifies this effect through complementary regulatory pathways^39, 40^.

The combination also demonstrated activity against CD34+ *KMT2A*-r leukemic stem and progenitor cells. Menin inhibition alone reduced colony forming potential, whereas selinexor in addition further amplified the potential while sparing normal hematopoietic stem cells even on long term exposure. The underlying mechanism is currently unknown. While detailed studies are needed to determine the mechanisms underlying this activity on stem and progenitor cells, it is interesting to note the combination of menin and XPO1 inhibitors is synergistic in multiple patient-derived CD34+ *KMT2A*-r and *NPM1*-m primary leukemic cells *in vitro*. Besides suppressing stemness, the combination treatment promptly activated monocytic differentiation by upregulating CD11b expression. The disruption of menin-KMT2A interaction in *KMT2A*-r, *NUP98*-r, and *NPM1*-m AML increased CD11b, which is consistent with our findings ^41–43^.

*In vivo* synergy between menin and XPO1 inhibitors in *KMT2A*-r models at reduced doses suggest the ability to maintain efficacy with improved tolerability, emphasizing the clinically relevant translational potential in AML. PDX models displayed survival beyond 6 months in the combination with a profound and rapid decrease in circulating hCD34+ leukemic cells, offering the potential to reduce the likelihood of acquisition of gatekeeper menin mutations. The combination was also very effective in suppressing leukemia in an aggressive PDX model derived from an *NPM1*-m AML patient sample, where a quick and substantial decrease in circulating leukemia burden was evident. The sustained effect despite treatment cessation, suggests the combination targeting the leukemic stemness, offering the potential for deeper responses and increasing the likelihood of achieving MRD negativity. Also, the leukemia burden in extramedullary sites including liver, and lung of combination-treated mice, was significantly lower, suggesting the combination has broad systemic efficacy.

Beyond MI-menin pocket binding site somatic resistance mutations, there is functional evidence to target cyclin-dependent kinases (CDK4/6) to resensitize resistant *KMT2A*-r leukemic cells to continued menin inhibition^28^. CDK6 is critical in MLL fusion-driven myeloid leukemogenesis^44^. It is also a target gene of MEIS1 in AML cells^45^. We observed the combination of menin-XPO1 inhibition inducing consistent differential expression of key cell cycle regulators in transcriptomic and proteomic studies in a *KMT2A*-r model (Supplementary Figure S6). This observation is interesting since it could support the combination to increase the efficacy of MI monotherapy, suggesting that as combination regimens anchored by menin inhibitors such as ziftomenib continue to be explored, a companion drug with a broader functional intersection like XPO1 inhibitor could be synergistic in multiple settings.

## Conclusion

Overall, our data support the combined inhibition of menin and XPO1 as a compelling strategy capable of enhancing transcriptional repression of the KMT2A/NPM1 leukemogenic program, reducing progenitor activity, promoting differentiation, and possibly overcoming resistance mechanisms. These findings provide a strong rationale for advancing ziftomenib-selinexor combinations into clinical evaluation across *KMT2A*-r and *NPM1*-m AML.

## Contributions

SKB conceived and led the study, supervised all aspects of the project, designed and performed experiments, analyzed data, and wrote the manuscript. MHU, SD, YH, AA, VD, JA, and SB performed research and analyzed the data. YD designed and performed research, analyzed data, and wrote the manuscript. AD, JY, JB, LP, LK, FB, AA, JC and JPM contributed critical reagents/materials and edited the manuscript.

## Acknowledgment

Work in SKB’s lab is supported by NIH/NCI Cancer Center Support Grant P30CA022453. Although this study was not directly funded by the Foundation, SKB gratefully acknowledges the UCANCERVIVE Foundation for their generous support of his laboratory’s overall research program. Kura Oncology, Inc. provided funding for part of the *in vivo* studies and ziftomenib under a sponsored research collaboration. Karyopharm Therapeutics provided selinexor as research support. The Biobanking and Correlative Sciences Core is supported, in part, by NIH Center grant P30 CA022453 to the Karmanos Cancer Institute at Wayne State University and by the Dresner Foundation. Work in YD’s lab is supported by Uniformed Services University of the Health Sciences (USUHS) Grant RAMP310534. The views presented in this manuscript are those of the authors; no endorsement by USUHS or the Department of War has been given or should be inferred.

