## Supplementary Information for "Combined Menin and XPO1 inhibition drive synergistic antileukemic activity in *KMT2A*r and *NPM1*-m AML"

### Figure S1

#### Interactions

| Gene ID A | Gene name A | Gene ID B | Gene name B | Organism | Type | Source | Score ? |
|---|---|---|---|---|---|---|---|
| ENSG00000133895 | MEN1 | ENSG00000283094 | PTEN | H. sapiens | SSL | Slorth | 0.886437 |
| ENSG00000100387 | RBX1 | ENSG00000133895 | MEN1 | H. sapiens | SSL | Slorth | 0.886197 |
| ENSG00000049759 | NEDD4L | ENSG00000133895 | MEN1 | H. sapiens | SSL | Slorth | 0.885932 |
| ENSG00000082898 | XPO1 | ENSG00000133895 | MEN1 | H. sapiens | SSL | Slorth | 0.885585 |
| ENSG00000133895 | MEN1 | ENSG00000165392 | WRN | H. sapiens | SSL | Slorth | 0.885379 |

**Figure S2**

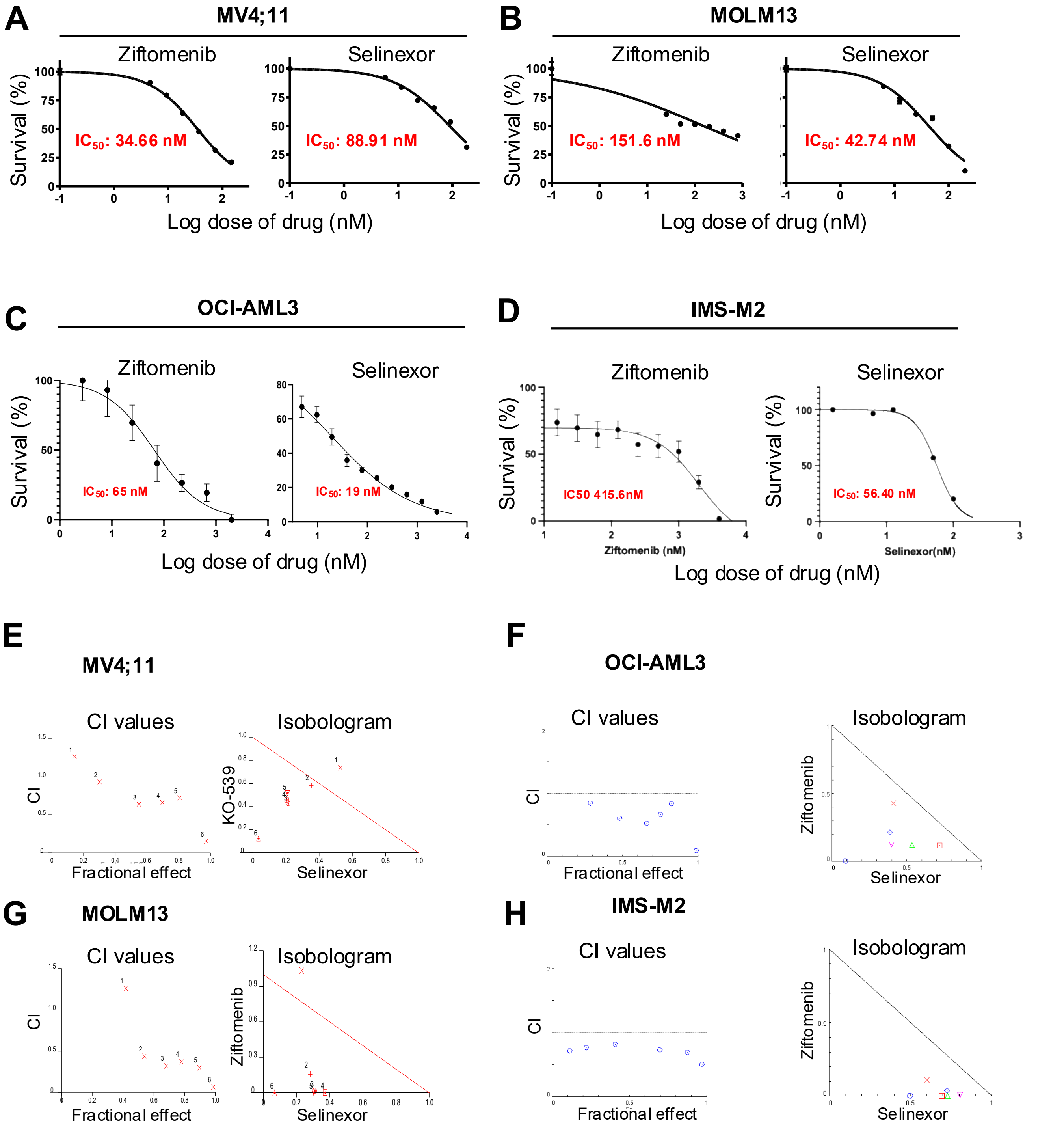


**I**

**Combination Index (CI)-IMS-M2**

| Ziftomenib (nM) | Selinexor (nM) | Effect | CI |
|---|---|---|---|
| 4000.0 | 160.0 | 0.97099 | 0.50124 |
| 2000.0 | 80.0 | 0.87764 | 0.69317 |
| 1000.0 | 40.0 | 0.69872 | 0.73145 |

**J**

**Combination Index (CI)-OCI-AML3**

| Ziftomenib (nM) | Selinexor (nM) | Effect | CI |
|---|---|---|---|
| 2000 | 160 | 0.82549 | 0.83803 |
| 1000 | 80 | 0.75247 | 0.65992 |
| 500 | 40 | 0.66361 | 0.52290 |

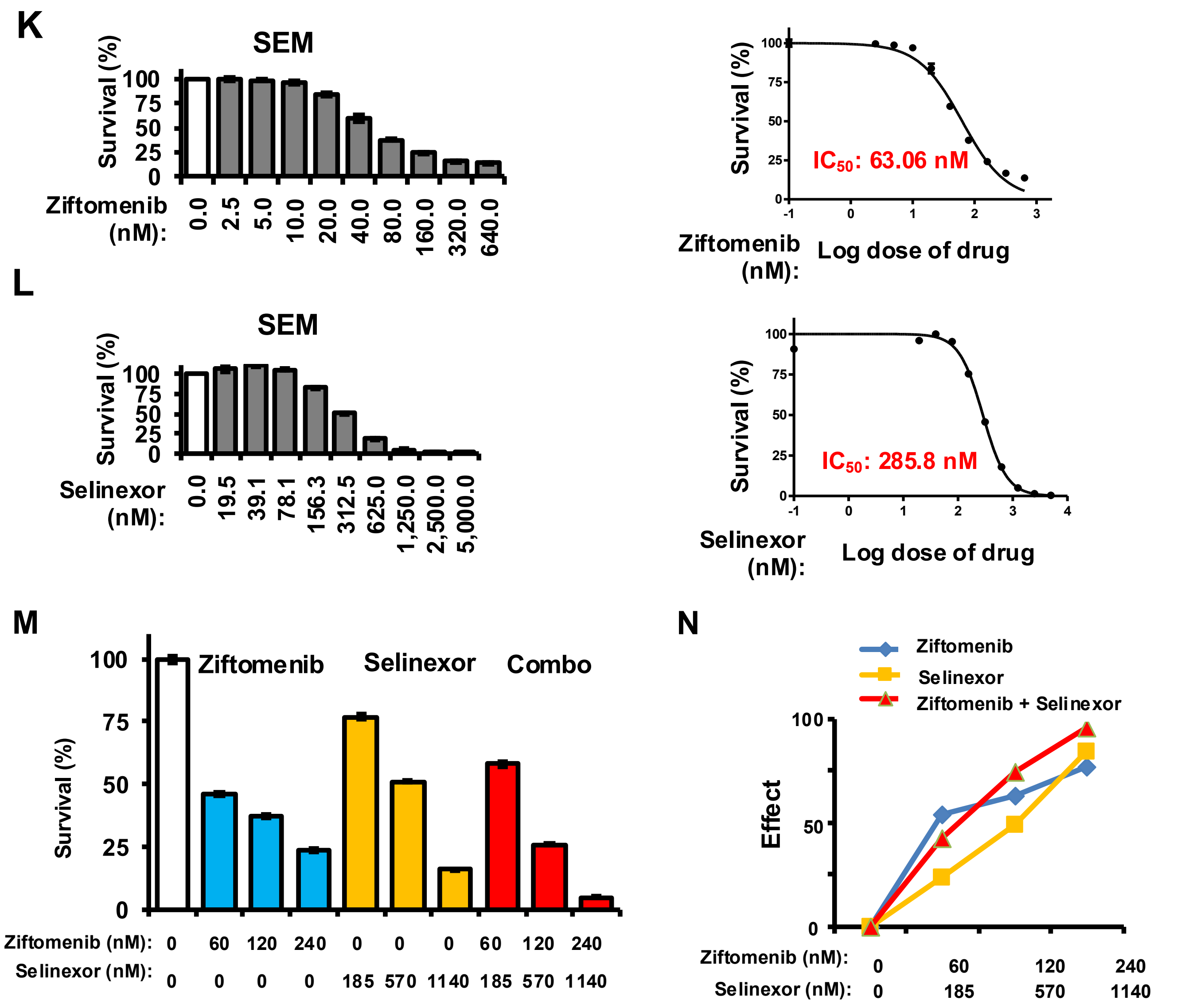


O

Combination Index (CI)-SEM

| Ziftomenib (nM) | Selinexor (nM) | Effect | CI |
|---|---|---|---|
| 60 | 285 | 0.419234 | 2.429 |
| 120 | 570 | 0.74289 | 1.194 |
| 240 | 1140 | 0.952881 | 0.580 |

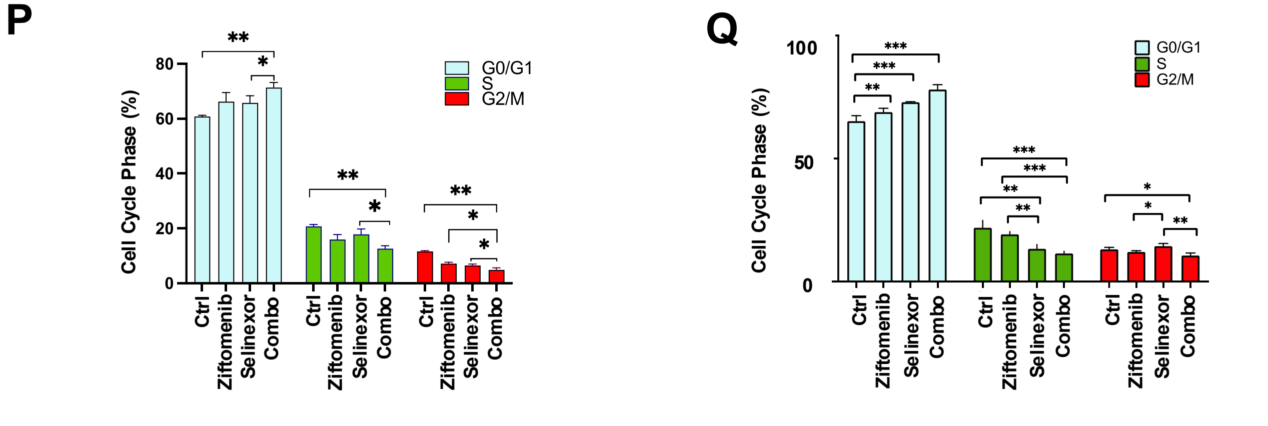

#### Figure S3

A

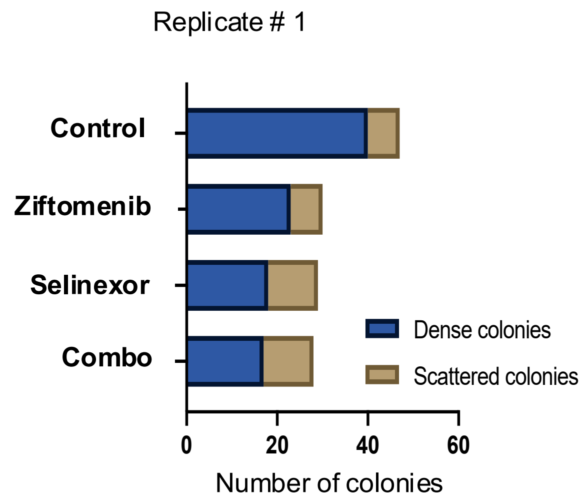
Replicate # 1
Control
Ziftomenib
Selinexor
Combo
Dense colonies
Scattered colonies
0
20
40
60
Number of colonies

B

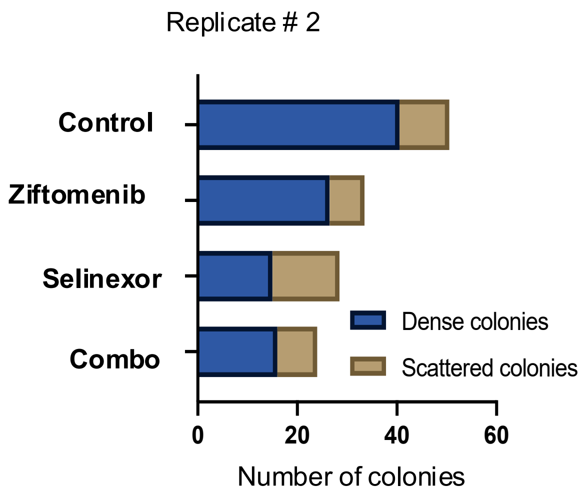
Replicate # 2
Control
Ziftomenib
Selinexor
Combo
Dense colonies
Scattered colonies
0
20
40
60
Number of colonies

Figure S4

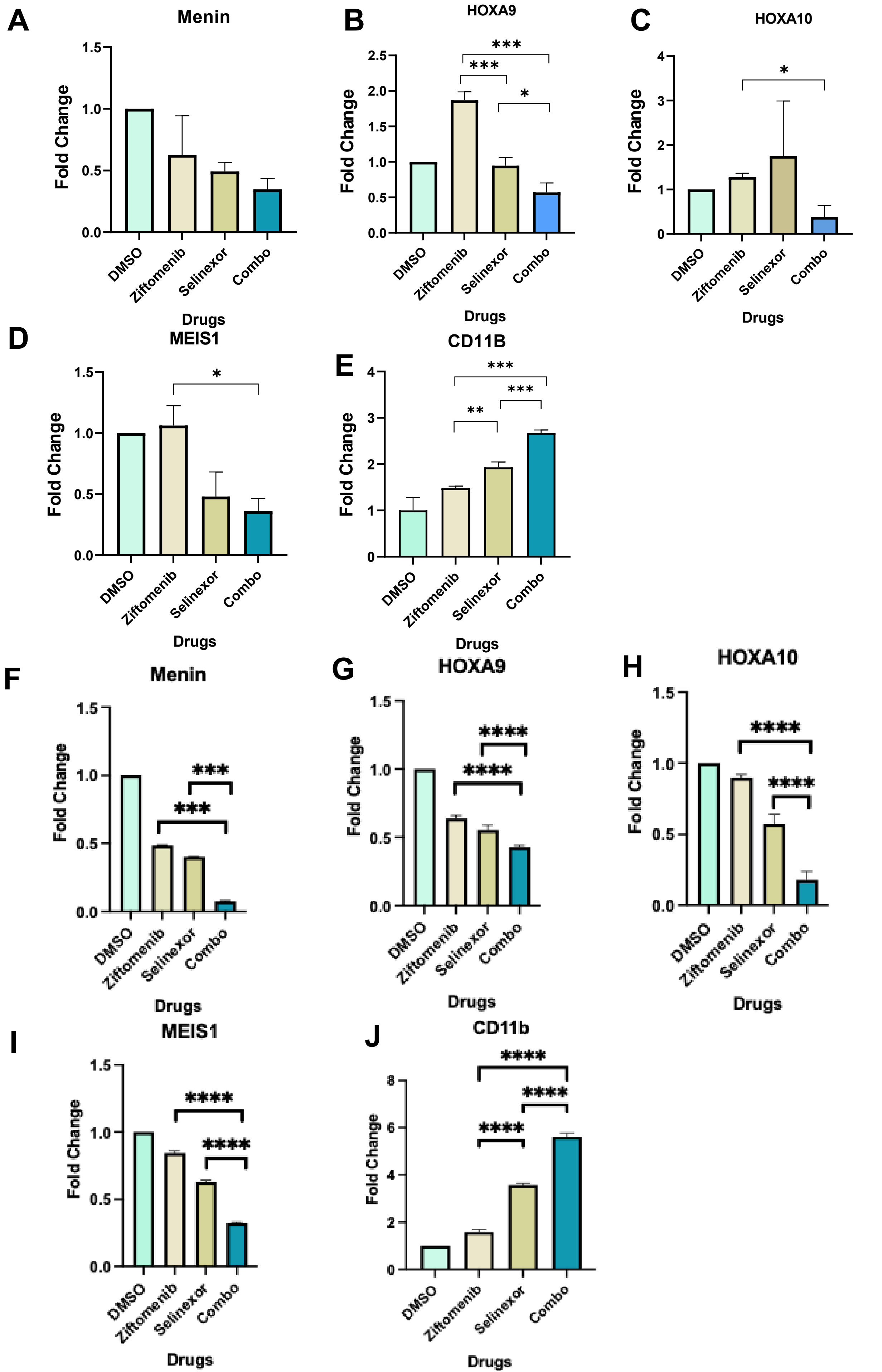

A
Menin
B
HOXA9
C
HOXA10
D
MEIS1
E
CD11B
F
Menin
G
HOXA9
H
HOXA10
I
MEIS1
J
CD11b
Fold Change
Drugs
DMSO
Ziftomenib
Selinexor
Combo

### Figure S5

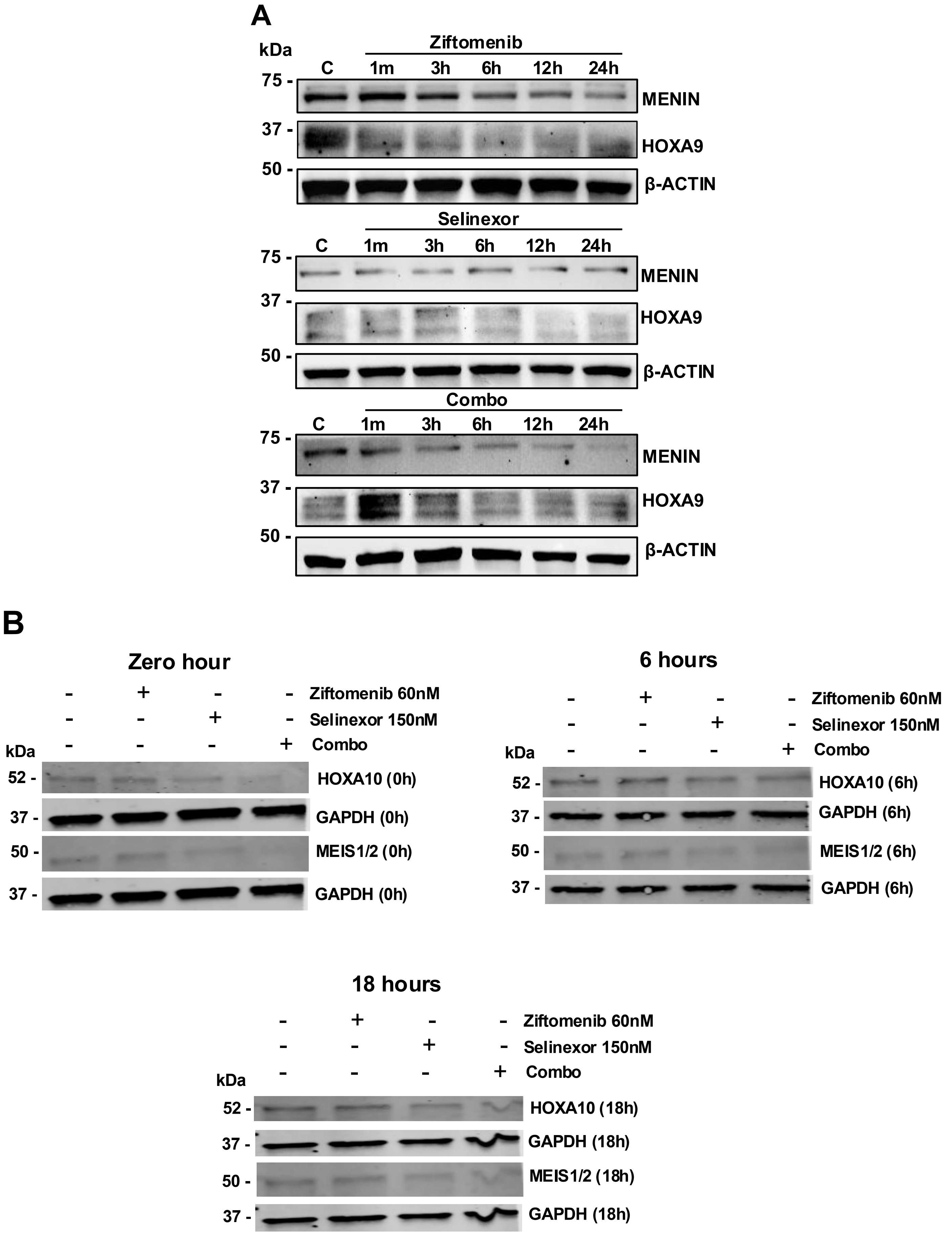
A
kDa
Ziftomenib
C 1m 3h 6h 12h 24h
75 -
MENIN
37 -
HOXA9
50 -
β-ACTIN
Selinexor
C 1m 3h 6h 12h 24h
75 -
MENIN
37 -
HOXA9
50 -
β-ACTIN
Combo
C 1m 3h 6h 12h 24h
75 -
MENIN
37 -
HOXA9
50 -
β-ACTIN
B
Zero hour
- + - - Ziftomenib 60nM
- - + - Selinexor 150nM
kDa - - - + Combo
52 - HOXA10 (0h)
37 - GAPDH (0h)
50 - MEIS1/2 (0h)
37 - GAPDH (0h)
6 hours
- + - - Ziftomenib 60nM
- - + - Selinexor 150nM
kDa - - - + Combo
52 - HOXA10 (6h)
37 - GAPDH (6h)
50 - MEIS1/2 (6h)
37 - GAPDH (6h)
18 hours
- + - - Ziftomenib 60nM
- - + - Selinexor 150nM
kDa - - - + Combo
52 - HOXA10 (18h)
37 - GAPDH (18h)
50 - MEIS1/2 (18h)
37 - GAPDH (18h)

C

#### Densitometry quantification of HOXA10 and MEIS1

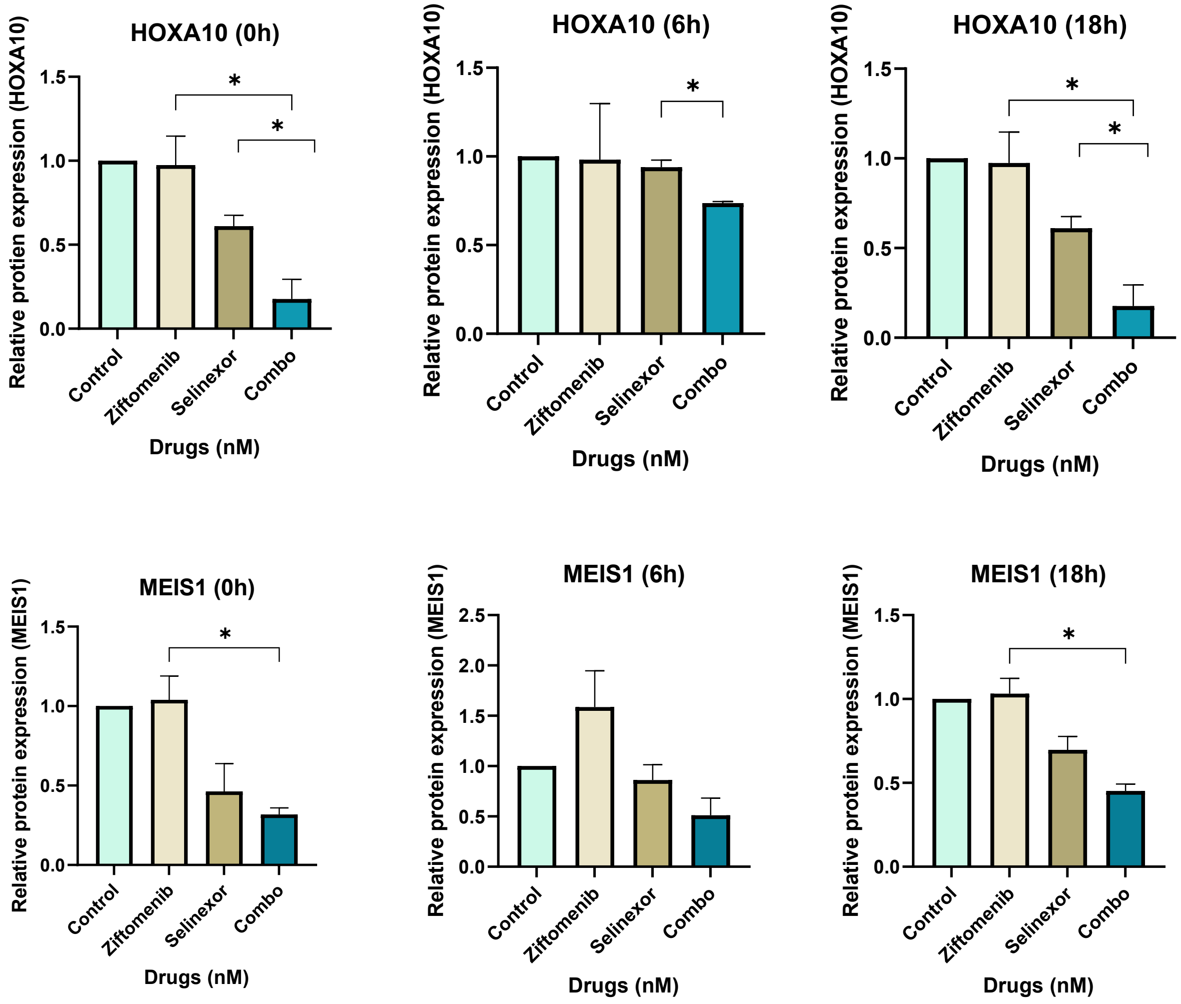

### Figure S6

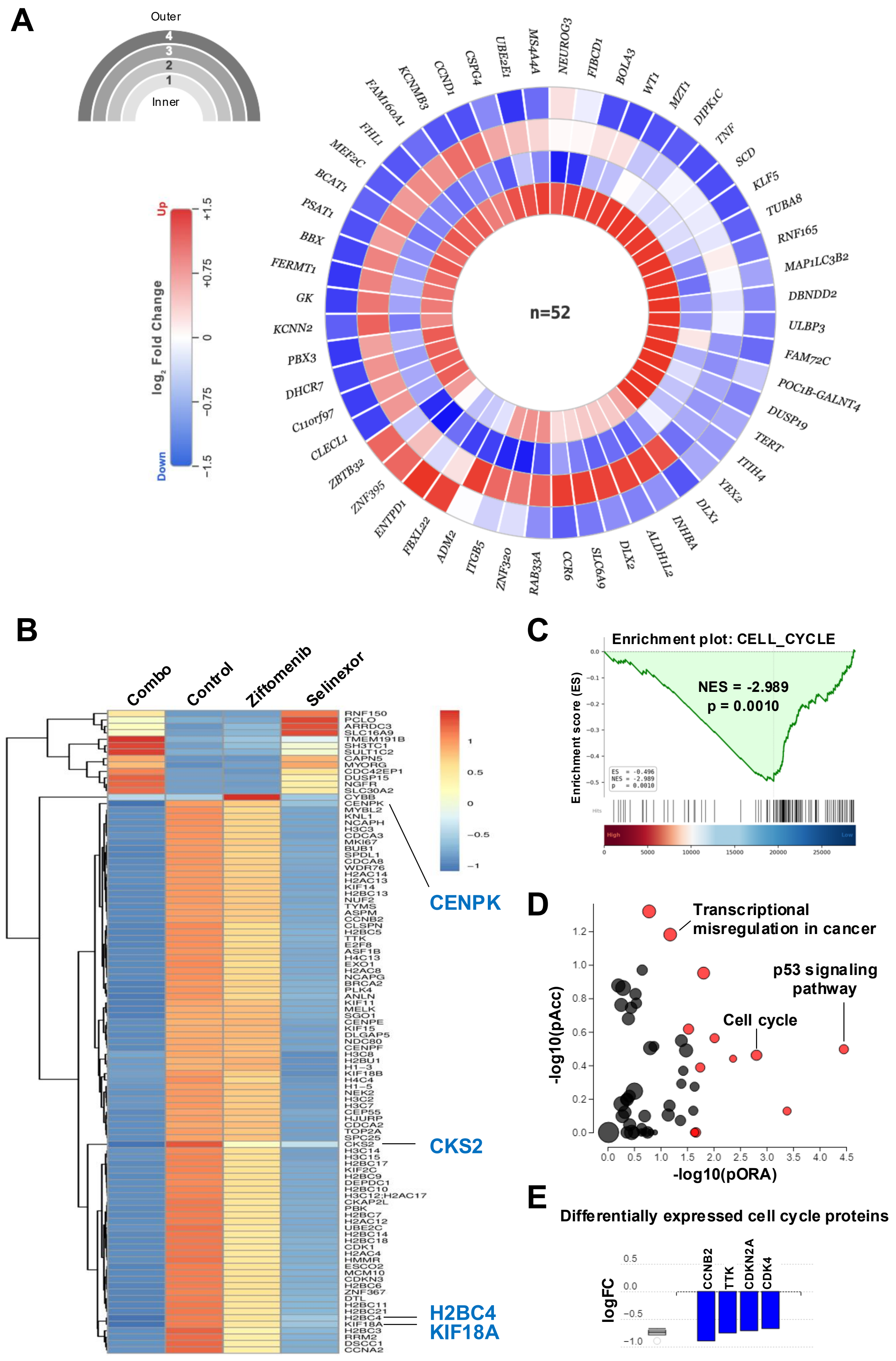
A
Outer
4
3
2
1
Inner
Up
log2 Fold Change
Down
+1.5
+0.75
0
−0.75
−1.5
n=52
B
Combo
Control
Ziftomenib
Selinexor
CENPK
CKS2
H2BC4
KIF18A
C
Enrichment plot: CELL_CYCLE
NES = -2.989
p = 0.0010
Enrichment score (ES)
D
Transcriptional misregulation in cancer
p53 signaling pathway
Cell cycle
-log10(pAcc)
-log10(pORA)
E
Differentially expressed cell cycle proteins
logFC
CCNB2
TTK
CDKN2A
CDK4

Figure S7

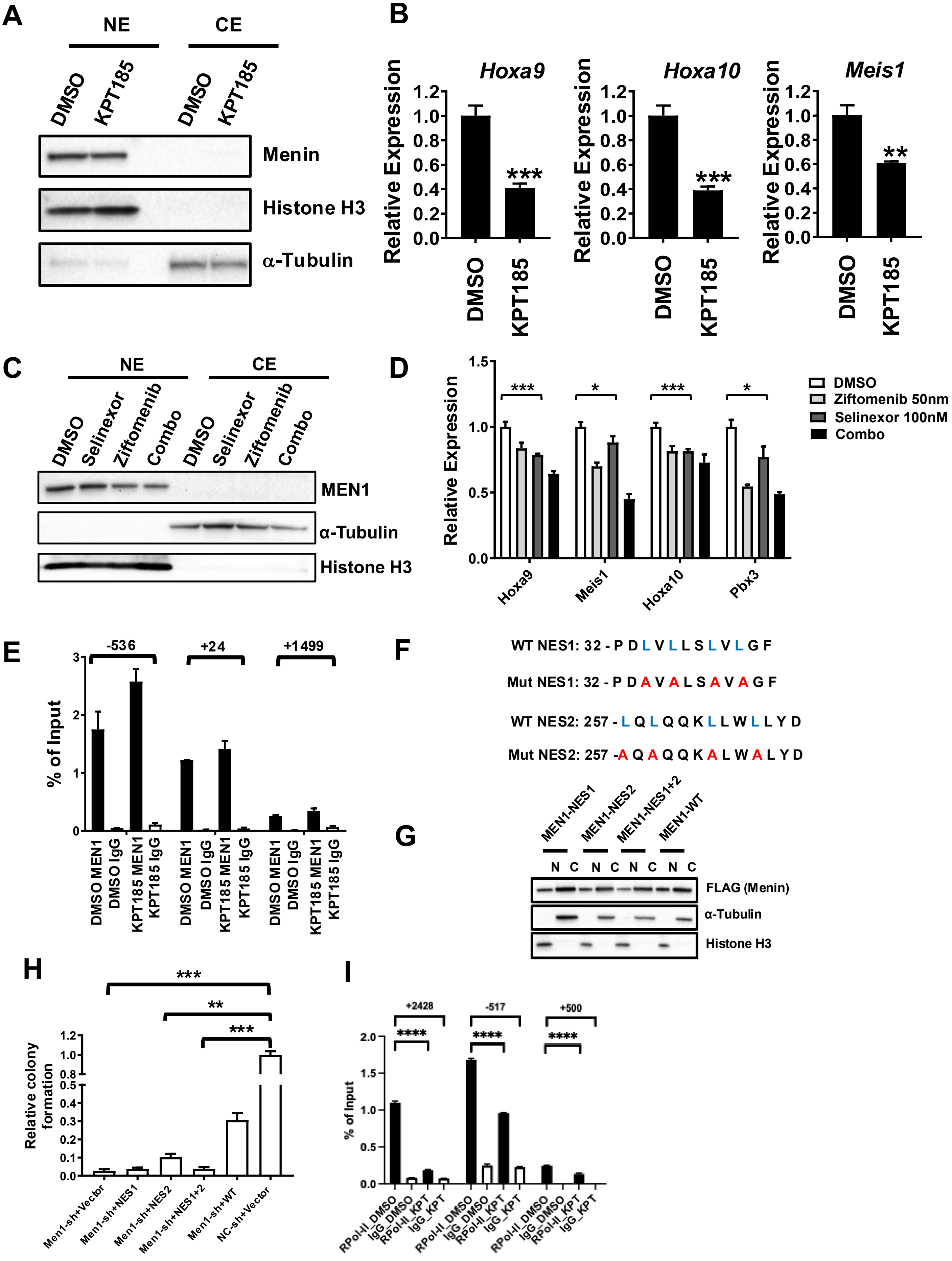

A
NE
CE
DMSO
KPT185
DMSO
KPT185
Menin
Histone H3
α-Tubulin
B
Hoxa9
Hoxa10
Meis1
Relative Expression
1.2
1.0
0.8
0.6
0.4
0.2
0.0
***
***
**
DMSO
KPT185
C
NE
CE
DMSO
Selinexor
Ziftomenib
Combo
MEN1
α-Tubulin
Histone H3
D
Relative Expression
1.5
1.0
0.5
0.0
***
*
***
*
DMSO
Ziftomenib 50nm
Selinexor 100nM
Combo
Hoxa9
Meis1
Hoxa10
Pbx3
E
-536
+24
+1499
% of Input
3
2
1
0
DMSO MEN1
DMSO IgG
KPT185 MEN1
KPT185 IgG
F
WT NES1: 32 - P D L V L L S L V L G F
Mut NES1: 32 - P D A V A L S A V A G F
WT NES2: 257 - L Q L Q Q K L L W L L Y D
Mut NES2: 257 -A Q A Q Q K A L W A L Y D
G
MEN1-NES1
MEN1-NES2
MEN1-NES1+2
MEN1-WT
N C N C N C N C
FLAG (Menin)
α-Tubulin
Histone H3
H
***
**
***
Relative colony formation
1.2
1.0
0.8
0.5
0.4
0.3
0.2
0.1
0.0
Men1-sh+Vector
Men1-sh+NES1
Men1-sh+NES2
Men1-sh+NES1+2
Men1-sh+WT
NC-sh+Vector
I
+2428
-517
+500
****
****
****
% of Input
2.0
1.5
1.0
0.5
0.0
RPol-II_DMSO
IgG_DMSO
RPol-II_KPT
IgG_KPT

#### Figure S8

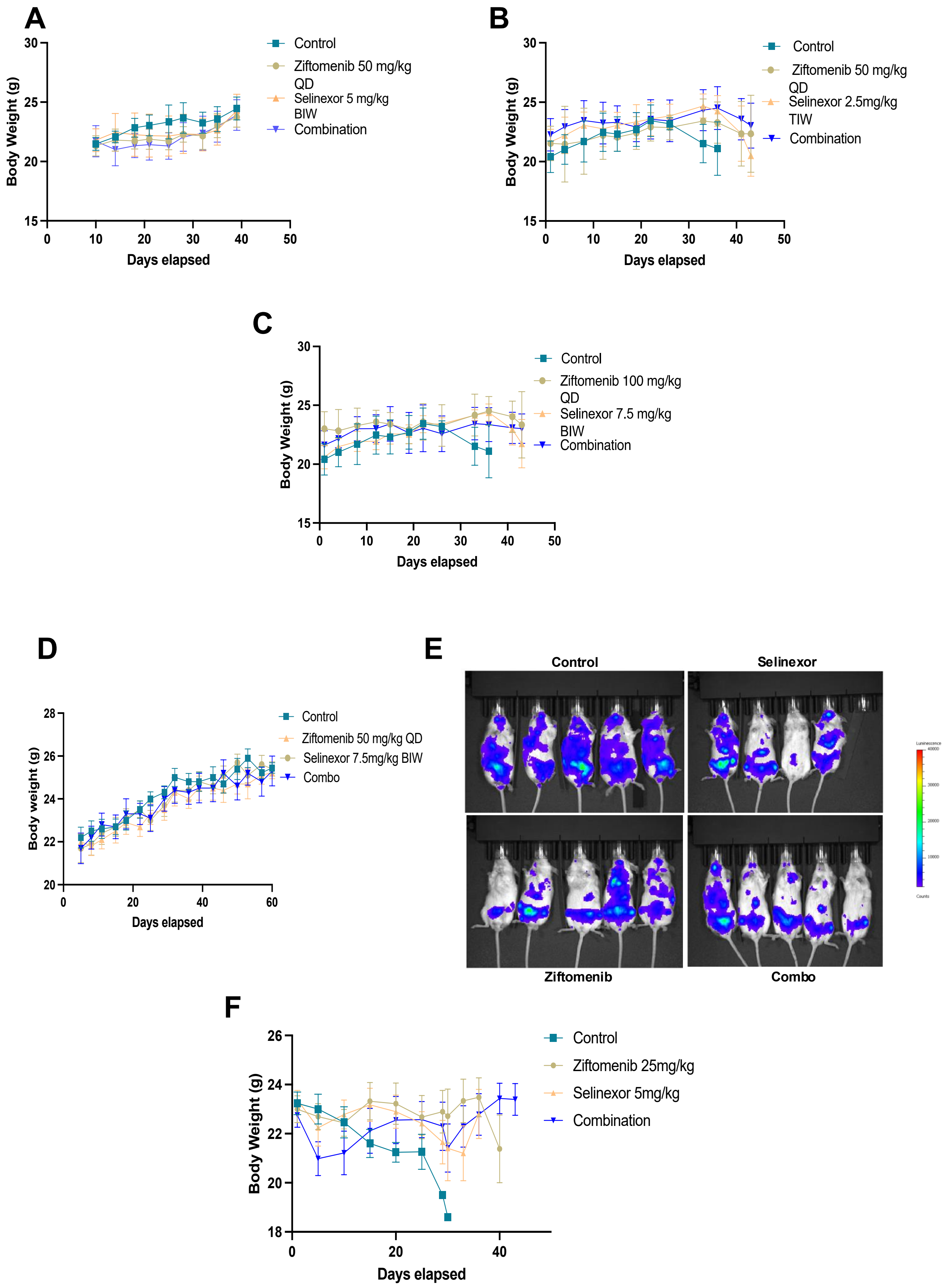
A
Body Weight (g)
Days elapsed
Control
Ziftomenib 50 mg/kg QD
Selinexor 5 mg/kg BIW
Combination
B
Body Weight (g)
Days elapsed
Control
Ziftomenib 50 mg/kg QD
Selinexor 2.5mg/kg TIW
Combination
C
Body Weight (g)
Days elapsed
Control
Ziftomenib 100 mg/kg QD
Selinexor 7.5 mg/kg BIW
Combination
D
Body weight (g)
Days elapsed
Control
Ziftomenib 50 mg/kg QD
Selinexor 7.5mg/kg BIW
Combo
E
Control
Selinexor
Ziftomenib
Combo
F
Body Weight (g)
Days elapsed
Control
Ziftomenib 25mg/kg
Selinexor 5mg/kg
Combination

### Table S1. List of primers and sequences used for RT-qPCR (human).

| Primers | Directions | Sequences (5'to 3') | References |
|---|---|---|---|
| GAPDH | F | CTCCTCCACCTTTGACGCTG | Shi et al., 2012 |
| | R | ACCACCCTGTTGCTGTAGCC | Eckelhart et al., 2011 |
| ACTIN | F | GGATGCAGAAGGAGATCACTG | Scheeren et al., 2008 |
| | R | CGATCCACACGGAGTACTTG | |
| 18S rRNA | F | CGGCTACCACATCCAAGGAA | Klimosch et al., 2013 |
| | R | GCTGGAATTACCGCGGCT | |
| STAT5B | F | CCGGGTAAACCATGGCTGTG | Yang et al., 2024 |
| | R | AGGAGCTGGGTGGCCTTAAT | |
| HOXA9 | F | ATGAGAGCGGCGGAGACAAG | Faaborg et al., 2021 |
| | R | GCACCGCTTTTTCCGAGTGG | |
| MEIS1 | F | CAGCCCATGGGAGGTTTCGT | Bhanvadia et al., 2018 |
| | R | GACCACCCCGGGCTACATAC | |
| HOXA10 | F | CTTCCGAGAGCAGCAAAGCC | Song et al., 2019 |
| | R | AGCCAGTTGGCTGCGTTTTC | |
| PBX3 | F | ATCGGCGACATCCTCCACCAG | Cheng et al., 2020 |
| | R | CTCATTAGCTGGGGATCGGGAG | |

### Table S2. List of primers and sequences used for RT-qPCR (mouse).

| Primers | Directions | Sequences (5'to 3') |
|---|---|---|
| Hoxa9 | F | CTCCTCCACCTTTGACGCTG |
| | R | ACCACCCTGTTGCTGTAGCC |
| Hoxa10 | F | GGATGCAGAAGGAGATCACTG |
| | R | CGATCCACACGGAGTACTTG |
| Meis1 | F | CGGCTACCACATCCAAGGAA |
| | R | GCTGGAATTACCGCGGCT |
| Pbx3 | F | CCGGGTAAACCATGGCTGTG |
| | R | AGGAGCTGGGTGGCCTTAAT |
| Actb | F | ATGAGAGCGGCGGAGACAAG |
| | R | GCACCGCTTTTTCCGAGTGG |

**Table S3. List of primers and sequences used for ChIP-qPCR for *HOXA9* promoter regions (human).**

| Primers | Sequence |
|---|---|
| *HOXA9(+2428)* Forward | 5’- GTGCCCACAAAGCTGTTTC-3’ |
| *HOXA9(+2428)* Reverse | 5’-GGGAGGAGTTGAAGGGAATG-3’ |
| *HOXA9(+517)* Forward | 5’-AAGTCGGAAACGACCAACAG-3’ |
| *HOXA9(+517)* Reverse | 5’-GCCAACCAAACACAACAGTC-3’ |
| *HOXA9(-500)* Forward | 5’-GCGAGGCAAACGAATCTGTT-3’ |
| *HOXA9(-500)* Reverse | 5’-CCAAATCGCATTGTCGCTCT-3’ |

**Table S4. List of primers and sequences used for ChIP-qPCR for *HOXA9* promoter regions (mouse).**

| Primers | Sequence |
|---|---|
| *Hoxa9(-536)* Forward | 5′- TGTCAGAGCGTTGGAAAGTG-3′ |
| *Hoxa9(-536)* Reverse | 5′- TGTGAATTTTGTGCCTTCCA-3′ |
| *Hoxa9(+24)* Forward | 5′- ACCAGAGCGGTTCATACAGG-3′ |
| *Hoxa9(+24)* Reverse | 5′- CAGACTGGAGATGGGGAAAA-3′ |
| *Hoxa9(+1499)* Forward | 5′- TGCCTGCTGCAGTGTATCAT-3′ |
| *Hoxa9(+1499)* Reverse | 5′- GAGCGGTTCAGGTTTAATGC-3′ |
| *Hoxa9(+1612)* Forward | 5’- GGTGCGCTCTCCTTCGC-3’ |
| *Hoxa9(+1612)* Reverse | 5’- GCATAGTCAGTCAGGGACAAAGTG-3’ |

#### Table S5. Combination Index (CI) of *KMT2A*-r primary leukemic samples

| Primary Patient Samples | Ziftomenib (nM) | Selinexor (nM) | Effect | CI |
|---|---|---|---|---|
| KCI-I | 1000<br>500<br>250 | 100<br>50<br>25 | 0.48267<br>0.36344<br>0.29209 | 0.68414<br>0.78183<br>0.70335 |
| KCI-II | 1000<br>500<br>250 | 150<br>75<br>37 | 0.57320<br>0.59531<br>0.51281 | 0.49262<br>0.21725<br>0.17410 |
| KCI-III | 1000<br>125<br>62.5 | 100<br>12.5<br>6.25 | 0.65075<br>0.36888<br>0.30623 | 0.58622<br>0.73140<br>0.63841 |
| KCI-IV | 1000<br>500<br>125 | 150<br>75<br>18.75 | 0.87332<br>0.82221<br>0.75300 | 0.15495<br>0.15146<br>0.12842 |
| KCI-V | 1000<br>500<br>250 | 150<br>75<br>37 | 0.59772<br>0.48959<br>0.32524 | 0.53675<br>0.47235<br>0.58009 |
| KCI-VI | 1000<br>500.0<br>250 | 150<br>75<br>37.5 | 0.76597<br>0.70604<br>0.56949 | 0.50069<br>0.42929<br>0.60721 |
| KCI-VII | 1000<br>500<br>250 | 150<br>75<br>37.5 | 0.63951<br>0.48227<br>0.33596 | 0.74455<br>0.64496<br>0.54580 |
| KCI-VIII | 1000<br>500 | 150<br>75 | 0.32810<br>0.19539 | 1.18316<br>0.83296 |
| KCI-IX | 1000<br>500<br>250 | 150<br>75<br>37.5 | 0.79545<br>0.70125<br>0.61975 | 0.13173<br>0.24258<br>0.31200 |

### Table S6. Combination Index (CI) of *NPM1*-m primary leukemic samples

| Primary Patient Samples | Ziftomenib (nM) | Selinexor (nM) | Effect | CI |
|---|---|---|---|---|
| NPM-I | 2000<br>1000<br>500 | 200<br>100<br>50 | 0.66787<br>0.51051<br>0.35039 | 0.28434<br>0.49038<br>0.85192 |
| NPM-II | 2000<br>250<br>62.5 | 200<br>25<br>6.25 | 0.82901<br>0.34504<br>0.24198 | 0.64878<br>0.95634<br>0.43218 |
| NPM-III | 2000<br>1000<br>500 | 200<br>100<br>50 | 0.80567<br>0.73987<br>0.59586 | 0.38499<br>0.30577<br>0.40935 |
| NPM-IV | 2000<br>1000<br>500 | 200<br>100<br>50 | 0.65242<br>0.49634<br>0.39191 | 0.27374<br>0.71277<br>4.78592 |
| NPM-V | 2000<br>1000<br>500 | 100<br>50<br>25 | 0.56037<br>0.41558<br>0.41727 | 0.93001<br>1.17843<br>0.58230 |

**Table S7. Cytogenetics, translocations and mutations of the *KMT2A*-r primary leukemic samples**

| KMT2A-r sample | Cytogenetics at sample time point | Translocations | Associated Mutations |
|---|---|---|---|
| KCI-I | 46,XX,t(1;9;11)(q12;p22;q23)/47, idem,+11 | MLL(KMT2A)/11q23 in 86.5% of cells | None |
| KCI-II | 47+R+48, XY, der(1)t(1;1)(p13;q25)t(1;9)(q21;q13),-6,+8, t(11;19)(q23;p13.3), +r, +mar[cp19]/46,XY | MLL(KMT2A)/11q23 in 75% of cells | None |
| KCI-III | 46, XY, t(6;11)(q27;q23)/46,XY | MLL/11q23 gene in 87% of cells | ETV6, BCOR |
| KCI-IV | 47,XX,+X,t(4;11)(q21;q23),(p13)/46,XX | "MLL(KMT2A)/11q23 gene region in 93% of cells" | None |
| KCI-V | 46 XX, t(9,11)(p22;q23)/46, XX | MLL/11q23 36% of cell | None |
| KCI-VI | 46 XX, t(9,11)(p22;q23)[12]/46, XX | MLL/11q23 gene in 88.5% cells | FLT3-ITD, STAG2 |
| KCI-VII | 48 XX, -7, +8, +8, +8, t(11,17)(q23;q25) | MLL/11q23 92% of cell | NRAS |
| KCI-VIII | 46 XY | MLL/11q23 87% of cell | IDH2, IDH2 |
| KCI-IX | 46,XY,t(9,11)(p22;q23) | MLL/11q23 86% of cell | None |

**Table S8. Cytogenetics, translocations and mutations of the *NPM1*-m primary leukemic samples**

| NPM-m sample | Cytogenetics at sample time point | Translocations | Associated Mutations |
|---|---|---|---|
| NPM-I | 46,XX,t(1;9;11)(q12;p22;q23)/47, idem,+11 | MLL(KMT2A)/11q23 in 86.5% of cells | None |
| NPM-II | 47+R+48, XY, der(1)t(1;1)(p13;q25)t(1;9)(q21;q13),-6,+8, t(11;19)(q23;p13.3), +r, +mar[cp19]/46,XY | MLL(KMT2A)/11q23 in 75% of cells | None |
| NPM-III | 46, XY, t(6;11)(q27;q23)/46,XY | MLL/11q23 gene in 87% of cells | ETV6, BCOR |
| NPM-IV | 47,XX,+X,t(4;11)(q21;q23),(p13)/46,XX | "MLL(KMT2A)/11q23 gene region in 93% of cells" | None |
| NPM-V | 46 XX, t(9,11)(p22;q23)/46, XX | MLL/11q23 36% of cell | None |
