## Supplementary Figure Legends for "Combined Menin and XPO1 inhibition drive synergistic antileukemic activity in *KMT2A*r and *NPM1*-m AML"

**Supplemental File**

**Supplementary Figure S1. Determination of synthetic lethal score (out of 1) using network topology computational systems approach** **generated with Slorth (http://slorth.biochem.sussex.ac.uk).** Top synthetic lethal (SSL) interactions between menin (MEN1) and other genes for human. SSL score between MEN1 and XPO1 is highlighted with a red box.

**Supplementary** **Figure S2. Effect of menin inhibitor and selinexor on the growth and deaths in AML cell lines.** (A-D) Growth inhibition of cells after KO-539 (ziftomenib) and selinexor treatments were determined using CellTiter Glo assay for MV4;11, MOLM13, OCI-AML3 and IMS-M2 cells. (E-J) Combination index (CI) values and normalized isobolograms for the MV4;11, MOLM13, OCI-AML3 and IMS-M2 cells treated with the combination of ziftomenib and selinexor. Most CI values are <1, demonstrating synergy between in ziftomenib and selinexor in all the cell lines. (K & L) Growth inhibition of SEM cells after ziftomenib and selinexor treatments respectively and their IC_50_ values (right panel). (M-O) Effect of ziftomenib and selinexor combination treatment on the growth of SEM cells. CI values and isobolograms were generated using Calcusyn 2.0 software. (P & Q) Cell cycle analysis across MV4;11 and OCI-AML3 cells respectively after 48 hours of treatment with ziftomenib, selinexor, and combination.

**Supplementary Figure S3.** (A &B) Quantification of colony numbers in primary KMT2A-rearranged AML progenitor cells after 14 days of treatment with control, ziftomenib, selinexor, or the combination, with colonies scored based on their morphology (dense vs. scattered). A and B indicate replicate 1 and 2 of primary KMT2A-rearranged AML progenitor cells.

**Supplementary Figure S4.** (A-J) Densitometric quantification of protein expressions (Menin, HOXA9, HOXA10, MEIS1, and CD11b) after 24h of treatment of ziftomenib, selinexor and combination (Immunoblots in Figure 3C&D).

**Supplementary Figure S5.** Effect of menin inhibitors and selinexor combination on menin associated signaling molecules in MV4;11 cells (A) Western blot for protein expression of Menin and HOXA9 in MV4;11 as determined in different time intervals (1m, 3m, 6h, 12h and 24h) of treatment. (B) Expression of menin and its downstream target proteins including HOXA10, MEIS1 at 0h, 6h and 18h time intervals. (C) Densitometric quantification of HOXA10 and MEIS1 protein expression at 0h, 6h and 18h of treatment.

**Supplemental Figure S6.** (A) Circular heatmap depicting significantly altered genes in control, ziftomenib, selinexor and combination treatment groups in MV4;11 cells. Circular heatmap displaying a curated set of genes (n=52) significantly altered by ziftomenib treatment, including canonical menin/KMT2A targets. Concentric rings represent expression changes across four conditions (inner to outer: Control, Ziftomenib, Selinexor, and Combination), with color intensity reflecting log₂ fold change (range −1.5 to +1.5). Significance was determined using DESeq2 with FDR < 0.05. (B) Heatmap showing differentially expressed genes in control, ziftomenib, selinexor and combination treatment groups in MV4;11 cells including CENPK, CKS2, KIF18A, and H2BC4. (C) Gene sets enrichment analysis (GSEA) of cell cycle in the combination treatment group. (D) The combination treatment exhibited significant alterations in key pathways compared to the control. The x-axis denotes over-representation (pORA), while the y-axis represents total pathway accumulation (pAcc). Each data point corresponds to a specific pathway, with the dot size proportional to the significance of the pathway. Significantly impacted pathways are highlighted in red, while non-significant ones are shown in grey. (E) Proteomic analysis revealing differential expressed cell cycle related proteins.

**Supplemental Figure S7.** (A) Western blotting analysis showing menin levels in nuclear (NE) and cytoplasmic (CE) extracts of *KMT2A/MLLT3*-immortalized mouse myeloid progenitor cells after treatment with KPT-185 (500 nM) or DMSO for 6 hours. (B)  Real-time RT-PCR analysis of *Hoxa9,* *Hoxa10* and *Meis1* mRNA levels in the same cells as in (A). Relative expression levels were calculated by normalizing to b-Actin mRNA levels in the same sample and DMSO-treated cells.  (C) Western blotting analysis showing Menin levels in nuclear and cytoplasmic extracts of *KMT2A/MLLT3*-immortalized myeloid progenitor cells after treatment with ziftomenib (50 nM), selinexor (100 nM), the combination of ziftomenib and selinexor, or DMSO for 6 hours. (D) Real-time RT-PCR analysis of *Hoxa9,* *Hoxa10*, *Meis1* and Pbx3 mRNA levels in the same cells as in (C). (E) ChIP-qPCR analysis of indicated *Hoxa9* regions using an anti-KMT2A-N antibody or control IgG in *KMT2A/MLLT3*-immortalized myeloid progenitor cells after treatment with KPT-185 (500 nM) or DMSO for 6 hours. (F) Alignment of mouse wild-type and mutant MEN1 NES1 and NES2 sequences. Indicated leucine residues (blue) were replaced with alanine residues (red). (G) Western blotting analyses of nuclear and cytoplasmic fractions of HEK293T cells transiently expressing indicated 3xFLAG-tagged Menin or Menin mutants with specified NES mutations using indicated antibodies. (H) Relative colony formation by *KMT2A/MLLT3*-immortalized myeloid progenitors on IMDM methylcellulose medium in the presence of mouse SCF (50ng/ml) and IL-3 (10ng/ml) after transduction with indicated combinations of empty pMYs retrovirus (Vector) or pMYs retrovirus expressing Menin NES mutants (NES1, NES2, and NES1+2) and a lentiviral shRNA targeting 3’UTR of endogenous Men1 (Men1-sh) or a non-targeting shRNA (NC-sh). Colony numbers were normalized to the Men1-sh + Vector control. (I) ChIP-qPCR analysis of RNA polymerase II (RPol-II) occupancy at the *HOXA9* promoter (-517 bp) and downstream regions (+500 bp and +2428) in MV4;11 cells treated with selinexor (KPT).

**Supplementary Figure S8. Effect of menin inhibitor and selinexor combination in cell line-derived/patient-derived xenograft (CDX/PDX) models.** (A-C) *KMT2A*r CDX mice body weight in grams (mean ±SEM) for figure 5A-C. (D) *KMT2A*r PDX mice body weight in grams (mean ±SEM). (E) GFP-luciferase-expressing OCI-AML3 CDX mice treated with vehicle control, selinexor, ziftomenib, or the combination, with bioluminescent imaging demonstrating differences in leukemic burden as reflected by luminescent signal intensity. (F) *NPM1*-m PDX mice body weight in grams (mean ±SEM).
